# Computational modeling of the evolutionary transition from C3 to C4 photosynthesis

**DOI:** 10.1101/2022.01.20.477074

**Authors:** Armin Dadras, Sayed-Amir Marashi, Kaveh Kavousi, Ali-Mohammad Banaei-Moghaddam

**Affiliations:** Department of Biotechnology, College of Science, University of Tehran, Tehran, Iran; Laboratory of Complex Biological Systems and Bioinformatics (CBB), Institute of Biochemistry and Biophysics (IBB), University of Tehran, Tehran, Iran; Laboratory of Genomics and Epigenomics (LGE), Department of Biochemistry, Institute of Biochemistry and Biophysics (IBB), University of Tehran

## Abstract

C4 photosynthesis is an evolutionary adaptation that minimizes the adverse effects of the high photorespiration rate. Although it is widely accepted that the C4 plants are evolved from C3 ancestors, the knowledge about the details of this process is yet to be complete. One application of constraint-based metabolic network modeling is to simulate evolutionary trajectories that an organism endures under a certain selective pressure. However, this approach is barely used to predict the evolution of a complex trait in eukaryotes. Here, we utilized a genetic algorithm combined with a constraint-based metabolic network model of *Arabidopsis thaliana* to simulate the trajectories of C3 to C4 conversion under high photorespiration rate conditions combined with different environmental conditions. Our modeling predicted that the C3-C4 intermediates and C4 strategies are superior to C3 photosynthesis in these environmental conditions. Besides, resource scarcities drive different evolutionary trajectories toward the emergence of C4 photosynthesis.

**AUTHOR SUMMARY:** It is estimated that high photorespiration reduces C3 crops productivity up to 50% [1]. Carbon concentrating mechanisms have evolved in many plants to overcome adverse effects of high photorespiration rate in especial environments. C4 photosynthesis is one of these adaptations that separate carbon assimilation reactions spatially between mesophyll and bundle sheath cells of the leaves. It is reported that C4 photosynthesis is evolved more than 65 times in 19 different families of angiosperms independently from C3 ancestors. Therefore, it is hypothesized that the potential modules, at the level of metabolism, anatomy, and gene regulation, which are needed to perform C4 photosynthesis are present in C3 plants. Since C4 plants grow better than C3 plants in hot and dry environments, scientists are working on converting C3 crops such as rice to C4 plants. However, several challenges, including a lack of understanding of the evolutionary paths toward manifestation of C4 trait, have hampered the C3-to-C4 engineering ([2]). Here, we used a metabolic network model of *Arabidopsis thaliana* as a C3 model plant and attempted to understand the possible evolutionary events leading to emerging of C4 photosynthesis in various scenarios and examined the results.

## INTRODUCTION

Photosynthesis is a process that is used by plants, algae, and certain prokaryotes to harvest light energy for converting atmospheric CO_2_ into carbohydrates. Most important crops, including wheat and rice, use the so-called “C3” photosynthetic mechanism. In this particular type of photosynthesis, the first step of the carbon fixation, *i.e.*, the carboxylation reaction, is catalyzed by an enzyme called Rubisco. In this carbon assimilation process, one mole of CO_2_ reacts with a mole of ribulose-1,5-bisphosphate (RBP). This reaction ends in the production of two moles of 3-phosphoglycerate (3PG), which is a three-carbon metabolite and is a precursor for the production of larger sugars such as glucose. Nevertheless, Rubisco also catalyzes an oxygenation reaction that consumes a mole of O_2_ and a mole of RBP to produce a mole of 3PG and a mole of 2-phosphoglycolate (PGL). The latter molecule is known to have inhibitory effects on certain photosynthetic enzymes [3]. Reactions of the photorespiration pathway recycle this molecule and convert it into 3PG. However, this conversion requires the consumption of high energy metabolites, like ATP and NADPH. Furthermore, as a result of the release of CO_2_ by photorespiration, biomass production is considerably reduced [4].

C4 photosynthesis is a carbon concentrating mechanism which is evolved in some plants to circumvent the unfavorable effects of high photorespiration rate [5]. Most C4 plants exhibit the so-called “Kranz anatomy”, which is a particular arrangement of cells in the leaves of these plants [6]. In this structure, mesophyll cells are clustered in a ring around the leaf veins outside the bundle sheath cells [7]. Furthermore, Rubisco activity is absent in mesophyll cells [8]. Kranz anatomy enables an increased CO_2_ concentration in bundle sheath cells, which, in turn, increases the carboxylation-to-oxygenation activity ratio of Rubisco [9]. In C4 plants, CO_2_ is converted to bicarbonate through the activity of carbonic anhydrase in the cytosol of the mesophyll cells. In these cells, phosphoenolpyruvate carboxylase (PEPC) catalyzes the reaction of bicarbonate with phosphoenolpyruvate (PEP) to form oxaloacetate, which is a four-carbon compound [10]. Because O_2_ is not a substrate of PEPC, compared to C3 plants, C4 plants are less sensitive to the high pressure of O_2_ in the leaves and have higher carbon to biomass conversion rate [11, 12].

Phylogenetic analysis of different C4 plants suggests that they are independently evolved more than 65 times in angiosperm lineages from C3 ancestors [13, 14]. Interestingly, all enzymes which are active in C4 photosynthesis are virtually present in the metabolic network of C3 plants, although they are not necessarily localized in the same compartment as C4 plants, or they are regulated differently [8]. Maize, sorghum, and millet are among the agriculturally important C4 plants. Metabolic components of C3 and C4 photosynthesis are shown schematically in Fig 1A and Fig 1B, respectively.

**Figure 1.**
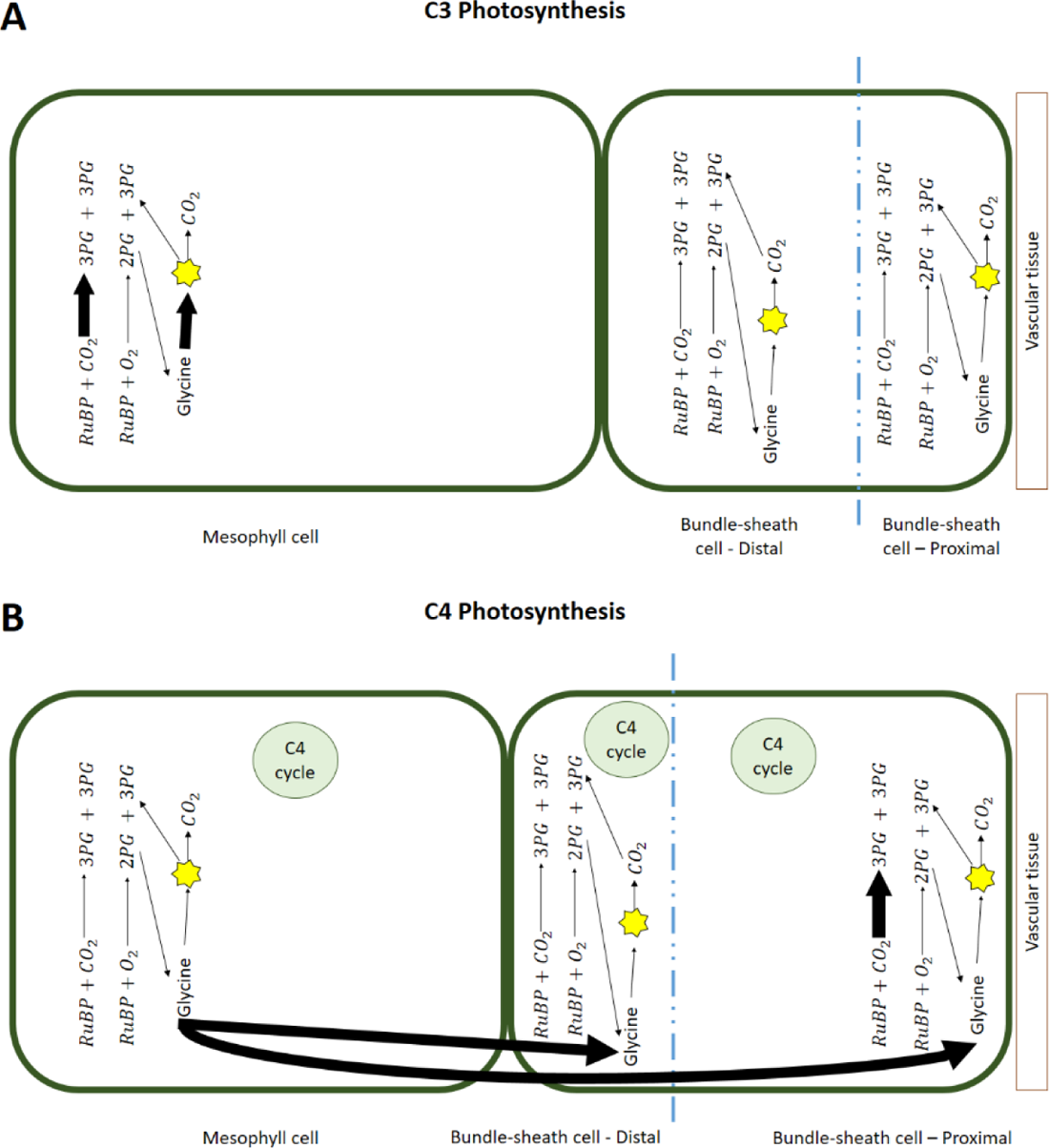
Schematic representation of metabolic components; in: A) a C3 plant; and B) a C4 plant. The thickness of the arrows symbolically indicates the activity of a pathway or reaction in a particular cell. The blue dotted lines represent the division of the bundle sheath cell into two proximal and distal parts. The yellow star schematically represents the reactions that convert glycine to 3-phosphoglycerate (3PG). The size of mesophyll cell and bundle sheath cell as well as the size of proximal and distal parts of bundle sheath cell are shown in proportion to the participation of these parts in the objective function.

While C4 plants are suggested to be the ideal crops in the era of global warming, efforts to create C4-like plants by genetically modifying C3 crops are yet to be successful [15]. This failure is partly because the evolutionary trajectories of the C3-to-C4 transition are not fully understood yet. The current understanding of the regulation and evolution of C4 photosynthesis is reviewed in detail elsewhere [16].

Constraint-based modeling (CBM) of metabolic networks has been previously used to simulate the reductive evolution of metabolism in obligate endosymbiotic bacteria [17, 18] and the adaptive evolution of metabolic networks by horizontal gene transfer [19–21]. In the present study, we utilized a genetic algorithm (GA) in combination with a constraint-based metabolic network model of *Arabidopsis thaliana* to simulate the evolution of C4 photosynthesis from C3 photosynthesis under high-rate photorespiration conditions. We considered five features of plants that have various forms in C3 compared to C4 plants. Our implementation was able to predict the optimal photosynthetic mechanism(s) and the evolutionary route(s) that lead to them under various environmental conditions. This observation suggests that under high photorespiration rate conditions, C4 photosynthesis (and depending on the imposed resource limitations, other photosynthetic strategies) can evolve from C3 photosynthesis. We present the graphical abstract of our study in Fig 2.

**Figure 2.**
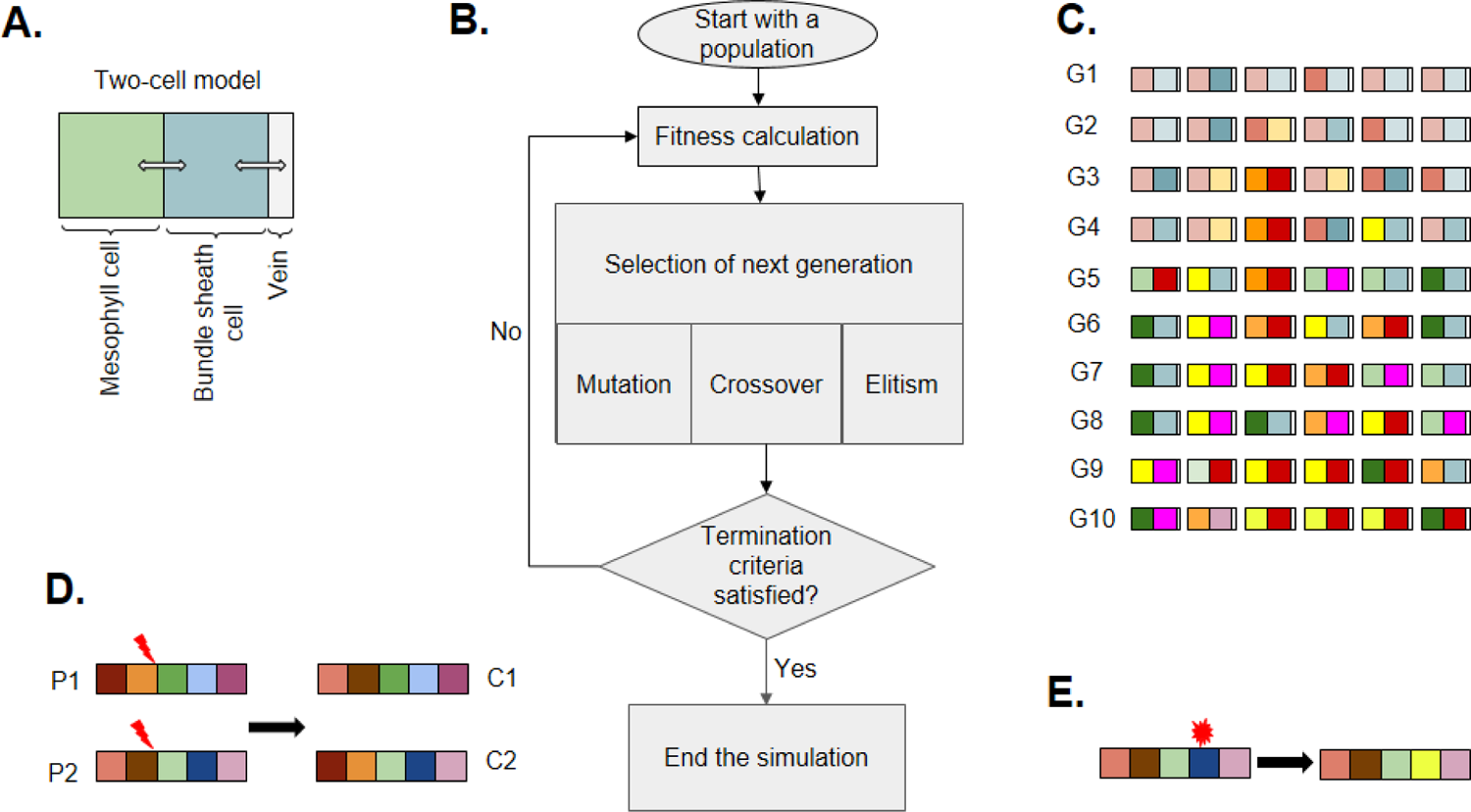
The schematic representation of the approach used in the current study. A) We create a two-cell model from an *A. thaliana* model. B) The flowchart shows how we implemented the genetic algorithm. One hundred instances of two-cell model created as the first generation. The fitness of each individual can be calculated from the fitness function. Using genetic algorithm operators such as crossover, mutation, and elitism, we produced next generation of population. We terminated our simulations after one hundred generations. C) Here, we show schematically how two-cell models (six arbitrary individuals) change over ten generations through genetic algorithms. D) In our implementation, we used one-point crossover to produce new individuals (children) from selected parents. E) In mutation, a specific parameter will change in the selected individual

## RESULTS

Since most C4 plants have the Kranz-type anatomy consisting of bundle sheath and mesophyll cells, we devised a two-cell model to allow simulations of C4 evolution. We modified the *A. thaliana* metabolic network model based on literature (see the Methods section). We combined two metabolic network models of *A. thaliana* to create a two-cell model, for simulating the interactions of mesophyll and bundle-sheath cells. A number of exchange reactions were also added to connect the mesophyll and bundle sheath parts of the model, and the boundary reactions of mesophyll and bundle sheath cells were modified accordingly. We describe the model creation steps in details in the S1 Appendix file.

### Comparison of two-cell model predictions for C3 and C4 states

To verify the relevance of our two-cell model, flux balance analysis (FBA) [22] was used to calculate different properties of a two-cell model which either utilizes C3 photosynthesis or C4 photosynthesis (assuming that all traits are set to the extreme levels). For FBA analysis, we used the constraints suggested for this model in previous studies [23, 24]. In addition to these constraints, those substances that enter the leaf cells through the vessels were allowed to merely enter the bundle sheath cells. As the objective function, we considered a linear combination of mesophyll cell biomass production, the distal part of the bundle sheath cell and the proximal part of the bundle sheath cell.

For five features of the FBA output, the two models were compared. These features are the maximum value of the objective function (that is, biomass production rate), the sum of fluxes for Rubisco reactions, the flux of photon input reaction for producing one unit of biomass, the flux of nitrogen input reactions for producing one unit of biomass, and the sum of the absolute values of the ATP-consuming fluxes. Table 1 summarizes the results of FBA for C3 and C4 models.

**Table 1.**
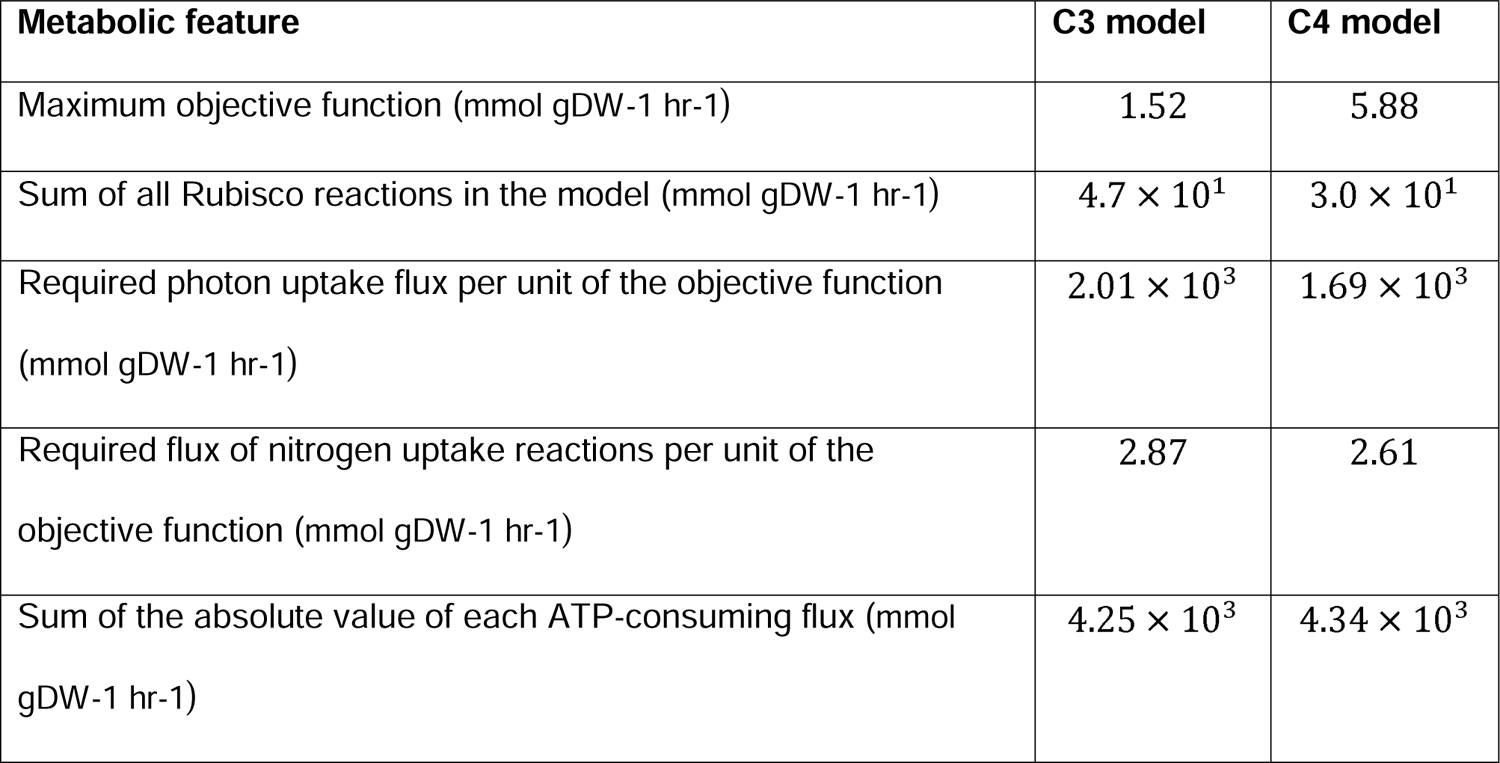
Evaluation of different properties of the two-cell models. FBA was applied to simulate the metabolic fluxes. Five distinguishing metabolic features are used to (qualitatively) compare the C3 and C4 models.

Our model predicted that C4 photosynthesis has: (i) higher maximum biomass production rate, (ii) needs less Rubisco flux, (iii) has higher efficiency in photon-to-biomass conversion rate, (iv) has higher nitrogen use efficiency, and (v) needs more ATP compared to C3 photosynthesis. These results suggest that the predictions of our C3 and C4 models are (at least) qualitatively acceptable.

### Modeling C4 evolution with genetic algorithm under resource limitation

In the next step, we combined the C3 model with a genetic algorithm (GA) to see if it is possible to simulate the evolution of C4 from C3 plants at high photorespiration rate. We addressed this question under four different scenarios. In the first scenario, there was no limitation on the resources. In each of the three other scenarios, the availability of a single resource (nitrogen, carbon, or water) was assumed to be restricted. We considered five key photosynthetic-related traits that were reported to be important in C4 evolution (Table 2). A “P-genotype” (photosynthetic genotype) was assigned to each individual as an ordered 5-tuple. For each P-genotype, one can assume a “P-phenotype” based on the biological knowledge about the characteristics of that certain P-genotype. For example, “11111” represents a C3 P-phenotype, while “44424” represents an individual with C4 P-phenotype. Additionally, for each P-genotype, one can reconstruct a corresponding metabolic model, for which a fitness value can be computed.

**Table 2.** Levels of traits in the GA and their photosynthetic equivalents. In S1 Appendix, the logic behind choosing each value is described in details.

|  |  |  |  |  |
| --- | --- | --- | --- | --- |
| Trait 1: The ratio of Rubisco reaction flux in mesophyll cell to this flux in bundle sheath cell |  |  |  |  |
| Trait values | 100 | 10 | 0.1 | 0.01 |
| Considered as | C3 | C3-C4 intermediate | C4 | C4 |
| Trait 2: The ratio of carboxylase to oxygenase activity of the Rubisco reaction in the proximal chloroplast of bundle sheath cell |  |  |  |  |
| Trait values | 1.139 | 1.937 | 3.530 | 4.130 |
| Considered as | C3 | C3-C4 intermediate | C3-C4 intermediate | C4 |
| Trait 3: Contribution ratio of the mesophyll, distal- and proximal-bundle sheath in the production of leaf biomass* |  |  |  |  |
| Trait values | 8.60 | 6.55 | 2.85 | 1.00 |
|  | 100 | 10 | 0.1 | 0.01 |
| Considered as | C3 | C3-C4 | C4 | C4 |
| Trait 4: The upper bound of selected C4 reactions |  |  |  |  |
| Trait values | 0 |  | 1000 |  |
| Considered as | C3 |  | C4 |  |
| Trait 5: The ratio of glycine decarboxylase reaction flux in the mesophyll cell to glycine transport flux (from mesophyll to bundle sheath) |  |  |  |  |
| Trait values | 100 | 10 | 0.1 | 0.01 |
| Considered as | C3 | C3-C4 intermediate | C3-C4 intermediate | C4 |

\* The second line indicates the contribution ratio of the distal bundle sheath reaction to the reaction in the proximal bundle sheath in total biomass production. If the second-line value is higher than one, the first-line indicates the flux ratio of the mesophyll reaction to the distal bundle sheath reaction contribution in biomass production otherwise determines the ratio of the mesophyll reaction to the proximal bundle sheath reaction contribution in biomass production. Also, the second-line indicates the ratio of reactions fluxes for the reactions in the distal to the proximal organelles of bundle sheath cell.

We initialized our first generation of GA with 70 individuals with C3 or C3-like P-genotypes. Then we introduced diversity in their P-genotypes, and chose the best individuals based on our selection method for creating the next generation (see the Method section and S1 Appendix file). For each scenario, we analyzed the population after 100 generations of “selection” and “creation of next generation” by genetic operators (crossover and mutation). In order to reduce the effect of chance, which is an inherent characteristic of GA, for each scenario the simulations were repeated 100 times.

Fig. 3 illustrates how the state of each P-trait evolves in the population (*i.e.*, a total of 70×100 individuals) over the 100 generations. In each panel, for a certain limitation scenario, the evolution of frequencies of individuals with different levels of a particular P-traits is plotted.

**Figure 3.**
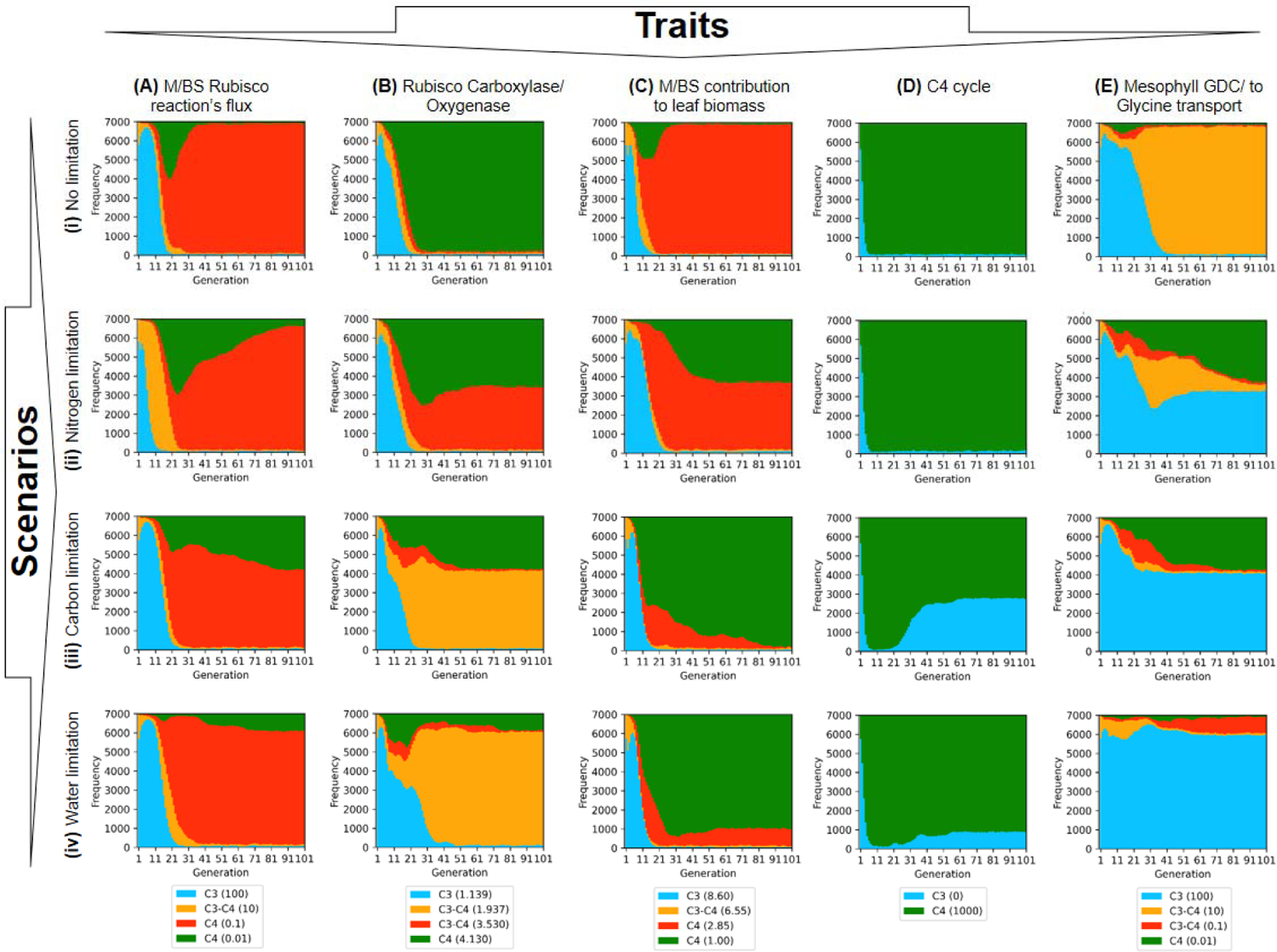
Evolution of individuals in each population with different states of a given P-trait in a particular condition. The state of each P-trait evolves in the population (*i.e.*, a total of 70×100 individuals) over the 100 generations. In each panel, for a certain limitation scenario, the evolution of frequencies of individuals with different levels of a particular P-traits is plotted.

When we investigated the Rubisco distribution between M and BS (trait 1, Fig 3. A_i_-A_iv_), we noticed that most of the individuals reached a C4 state in all scenarios. More specifically, in all scenarios, the majority of the individuals reached a state that Rubisco activity is ten times higher in BS compared to M. Since the current accepted evolutionary theory of C4 evolution suggests that Rubisco activity disappears from M cells, one might expect to observe more green (M/BS =0.01) individuals than red ones.

In the N-limited scenario (A_ii_), there is an interesting pattern in the M/BS Rubisco reaction flux: up to half of the population reaches the extreme C4 state (M/BS = 0.01) in the 31st generation, but then, the other C4 level (M/BS = 0.1) dominates the population.

Fig. 3 B_i_-B_iv_ represents the ratio of carboxylase to oxygenase Rubisco activity in the proximal chloroplast of bundle sheath cell. In no limitation scenario, the carboxylase-to-oxygenase activity of Rubisco in the proximal BS cell (C/O) reached the C4 state in most individuals (Fig. 3 B_i_). In the N-limited scenario (Fig. 3 B_ii_), we observed that only about half of the individuals reach the C4 state. On the other hand, in C- and W-limited scenarios, an intermediate state dominates the population. Besides, when water was limited, C4 state initially increases in the population, but later on, the frequency of C4 individuals deceases (Fig. 3B_iv_).

Based on Fig. 3 C_i_-C_iv_, we observed that the ratio of M to BS reached a C4 state in all four scenarios. One can see that in carbon or water limitation scenarios 1:1 ratio is favored, while in the other two scenarios, 2.85:1 is more favorable.

Figs. D_i_-D_iv_ shows the activity of C4 enzymes during the course of simulations. We observed that in the no-limitation and N-limited scenarios, C4 cycle enzymes rapidly reach their maximum activity, and then, stably remain in the population. In contrast, in the C- and W-limited scenario, after the initial activation of C4 cycle enzymes, some individuals revert to the initial state.

Fig. 3 E_i_-E_iv_ represents the formation of the glycine pump mechanism. In the no-limitation scenario, we see that a weak form of glycine pump mechanism is established, and subsequently, dominates the population (Fig. 3 E_i_). In this primitive form, for decarboxylation of every ten mole of glycine by GDC, one mole glycine is transported from M to BS. In contrast, in the other three scenarios, at least in half of the individuals, glycine pump mechanism is not evolved. In C- and N-limited scenarios, in about one-third of the individuals a fully functional glycine pump mechanism is evolved. In contrast, one can observe that in the W-limited scenario, most of the individuals maintained the C3 state for this trait, that is, only 1 in 100 molecule of glycine is transported to BS.

### Evolutionary trajectories that lead to C4 photosynthesis

It is already shown that the formation and the ecological distribution of C4 photosynthesis is presumably driven by a complex interaction of environmental conditions with pre-existing genomic and anatomical enablers [10, 25]. Therefore, to better understand the trajectories of C4 evolution, we investigated how evolution of each trait affects the evolution of other traits.

### The no-limitation scenario

We firstly focused on the population that was present in the last (100^th^) generation of simulation under the no-limitation scenario (*i.e*., 70×100 individuals). For each P-genotype that is present in the last generation, the relative frequency of that individual is shown in Fig. 4A.

**Figure 4.**
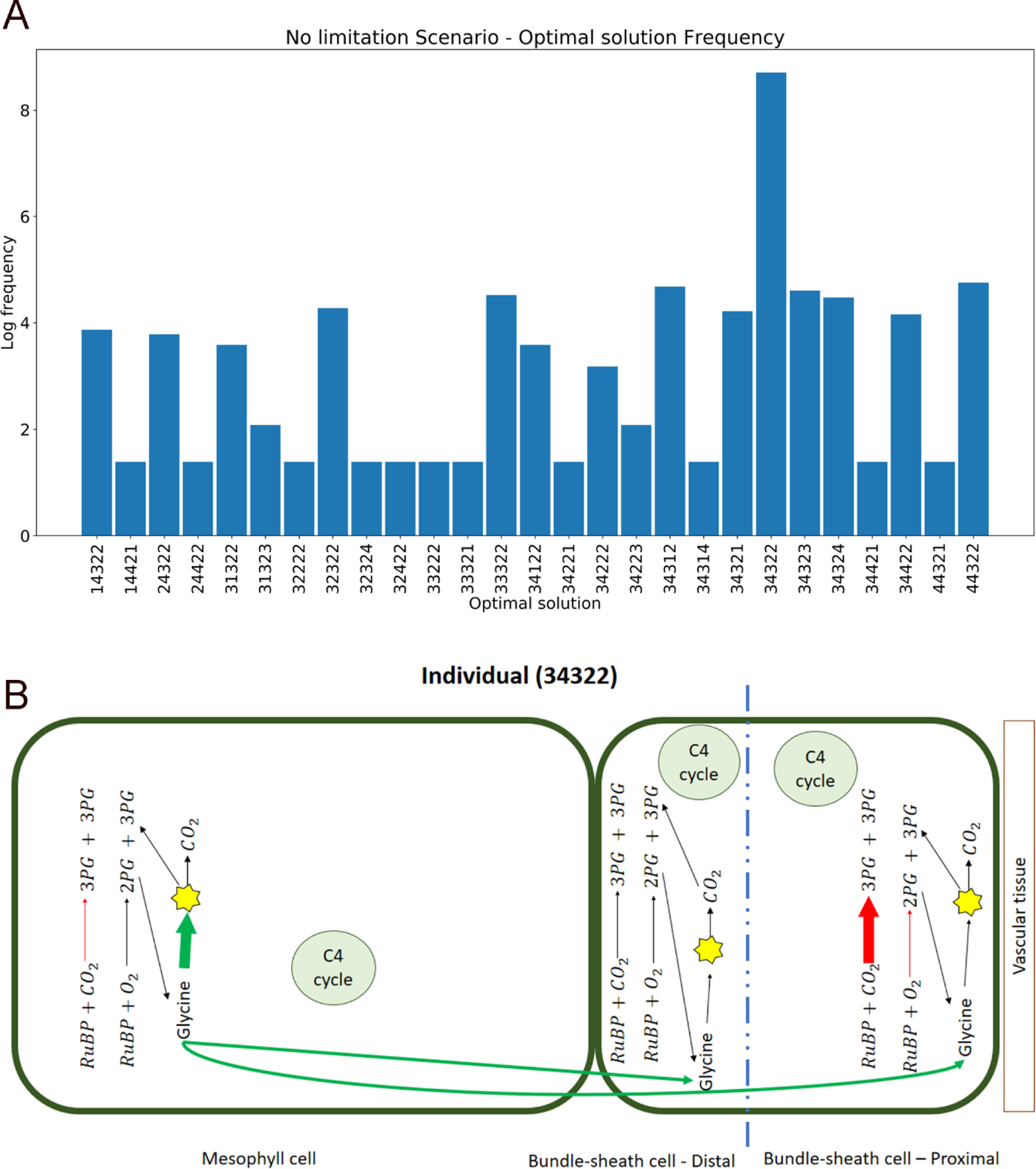
The “no limitation” scenario. A) The logarithm of frequency of selected individuals (solution) in the last (100^th^) generation; B) Schematic representation of traits in simulated metabolic pathways of the most frequent individual.

In the no-limitation scenario, according to Fig. 4A, P-genotype of the dominant individual was “34322”, which means that it has the C4 level for all traits except for the 5th trait (the ratio of glycine decarboxylase reaction flux in the mesophyll cell to glycine transport flux from mesophyll to bundle sheath). We presume that this individual utilizes a C4-like photosynthetic strategy. A schematic representation of this individual is shown in Fig. 4B.

To better understand the evolutionary trajectories which results in the selected individuals, we showed the most abundant individuals using a directed graph (Fig. 5). To simplify the figures, individuals of every ten generations were grouped and the most frequent individuals were shown in a single column. However, this led to removal of rare individuals, and therefore, our graph is not (necessarily) connected in certain scenarios. For example, the first column depicts the most frequent individuals in generations 1 to 10. When a specific photosynthetic mechanism emerged in a generation, we did not redraw it in the next generations even if it was present in the next one.

**Figure 5.**
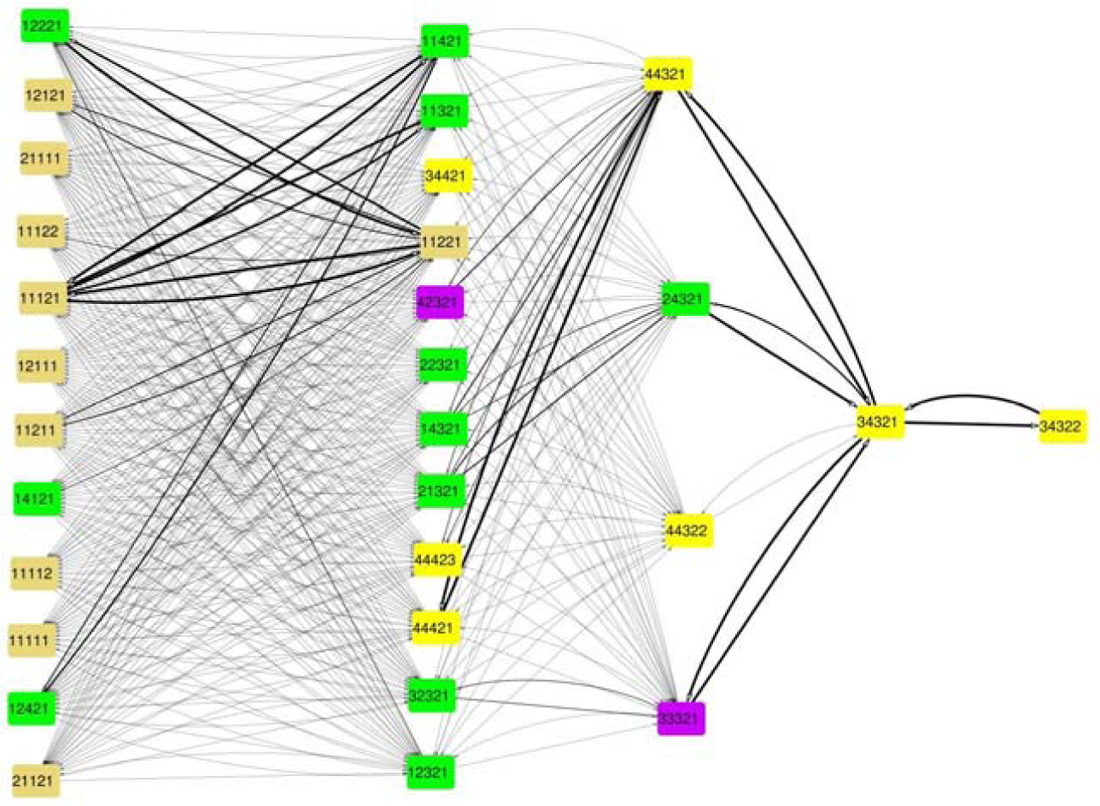
The trend of changes in population in the “no limitation” scenario. The pink, green, purple, and yellow circles are individuals with C3-like, C3-C4 intermediate, C4-like, C4 photosynthesis, respectively. Since we drew only individuals with high frequency in the population, this graph is not necessarily connected. Nodes represent the individuals, and the thickness of each arrow represents the “chance” to observe a transition from the first individual to the second one. The width of each edge is proportional to the log frequency transition from the source node to the destination node as explained in methods section.

### Scenario 2: N-limited scenario

The logarithm of relative frequency for each P-genotype that is present in the last generation’s individuals in N-limited scenario is plotted in Fig 6A, similar to previous Scenario.

**Figure 6.**
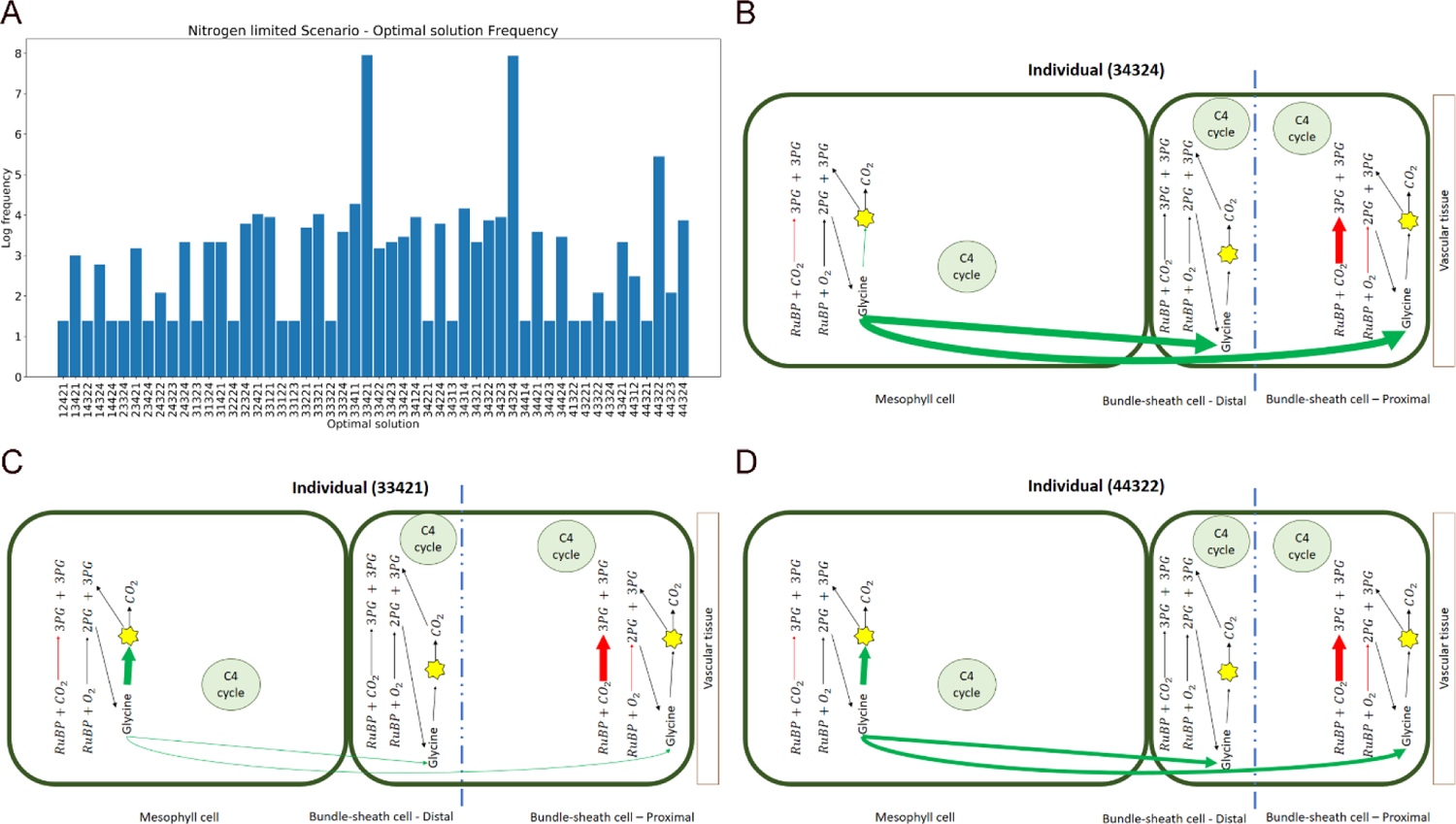
Nitrogen limitation scenario. A) The logarithm of frequency of selected individuals in the last (100^th^) generation; B, C, D) Schematic representation of traits in simulated metabolic pathways of the top three individuals. The thickness of the arrows symbolically indicates the activity of a pathway or reaction in a particular cell. The blue dotted line represents the division of the bundle sheath cell into two proximal and distal parts. The yellow star schematically represents the reactions that convert glycine to 3PG. The size of mesophyll cell and bundle sheath cell as well as the size of proximal and distal parts of bundle sheath cell are shown in proportion to the participation of these parts in the objective function.

Based on Fig 6A, in the N-limited scenario, two individuals, “33421” and “34324”, have the best performance among all, which are equivalent to a version of proto-Kranz and C4 strategy. We schematically represent the traits of these individuals in Fig 6B and Fig 6C, respectively. The third optimal photosynthetic strategy in this scenario is C4-like, “44322” and its traits are represented in Fig 6D.

As in the previous Sections, we showed the evolutionary trajectories of individuals using a directed graph for nitrogen limitation simulation in Fig. 7.

**Figure 7.**
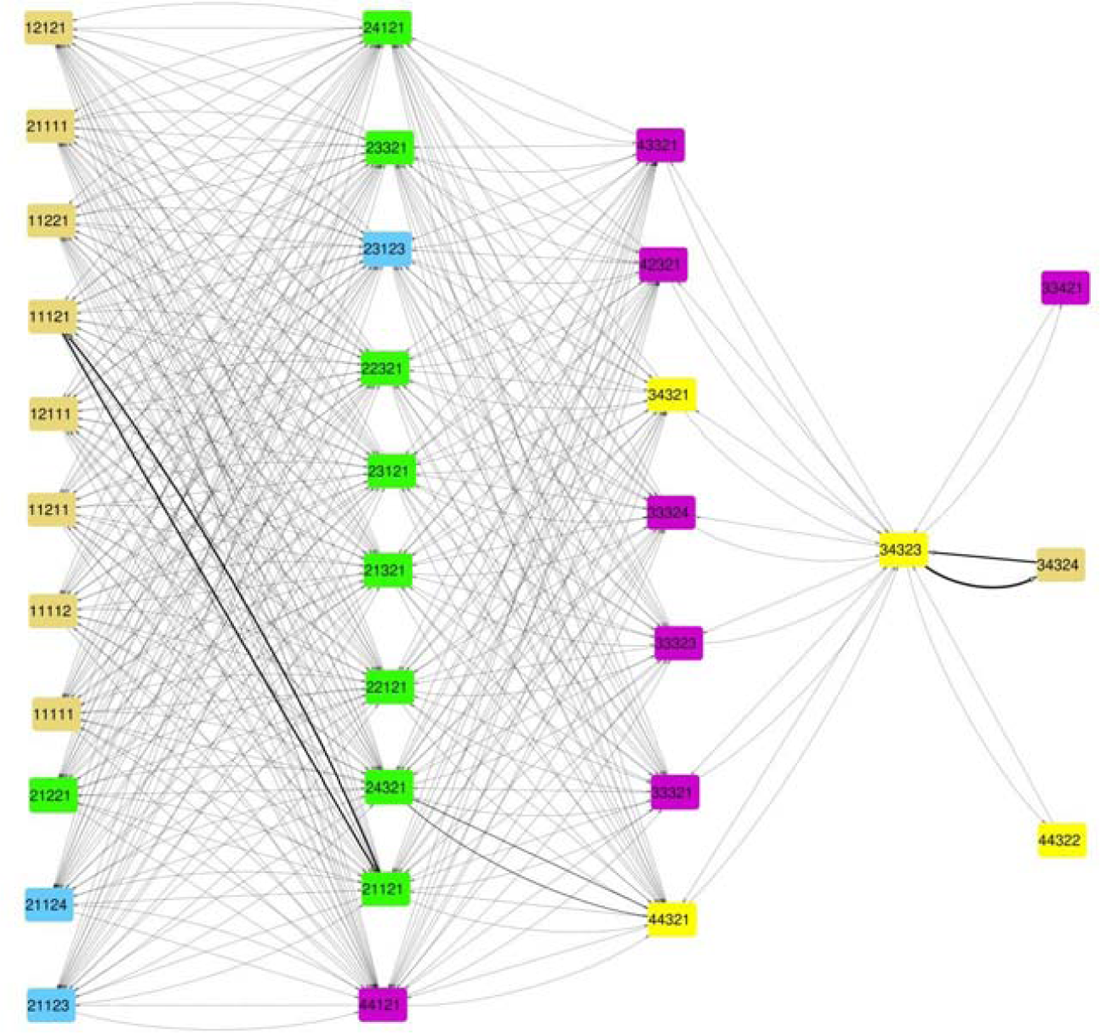
The trend of changes in population in nitrogen-limited scenario. The pink, green, cyan, purple, and yellow circles are individuals with C3-like, C3-C4 intermediate, C2, C4-like, C4 photosynthesis, respectively. Since we drew only individuals with high frequency in the population, this graph is not necessarily connected. Nodes represent the individuals, and the thickness of each arrow represents the “chance” to observe a transition from the first individual to the second one. The width of each edge is proportional to the log frequency transition from the source node to the destination node as explained in Methods section.

### Scenario 3: Carbon limitation

Similar to Scenario 2, we gathered the information of all individual that were present in the last (100^th^) generation of this scenario. The logarithm of frequencies of individuals that are present in the last generation, *i.e.*, the optimal solutions found for carbon limitation scenario, are plotted in Fig 8A.

**Figure 8.**
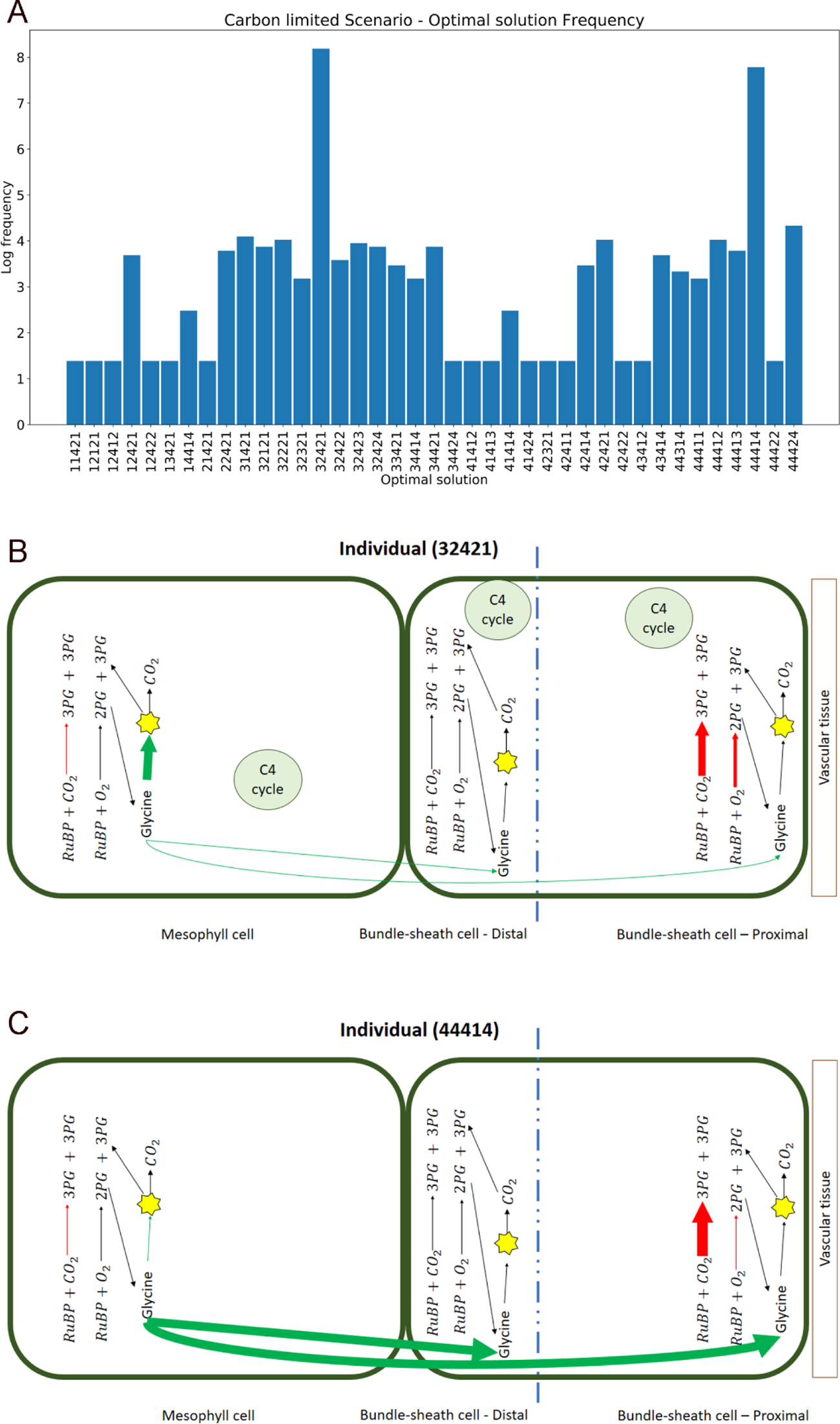
Carbon limitation scenario. A) The logarithm of frequency of optimal individuals in the last (100^th^) generation; B, C) Schematic representation of traits in simulated metabolic pathways of the top two individuals The thickness of the arrows symbolically indicates the activity of a pathway or reaction in a particular cell. The blue dotted line represents the division of the bundle sheath cell into two proximal and distal parts. The yellow star schematically represents the reactions that convert glycine to 3PG. The size of mesophyll cell and bundle sheath cell as well as the size of proximal and distal parts of bundle sheath cell are shown in proportion to the participation of these parts in the objective function.

According to Fig 8A, in the carbon scarcity scenario, the “32421” and “44414” individuals that are equivalent to C4-like and C2 photosynthesis, respectively, are the optimal photosynthetic strategies. The traits of these two models are schematically shown in Fig 8B and Fig 8C, respectively. Moreover, Fig 8A shows that in this scenario, the number of distinct optimal solutions is higher compared to the “no limitation” scenario. This is probably because carbon scarcity has reduced the authority of dominant strategies in the population and provided the opportunity for less-efficient strategies to stay present in the population.

Similar to Scenario 1, we showed the evolutionary trajectories of individuals using a directed graph for carbon limitation simulation in Fig. 9.

**Figure 9.**
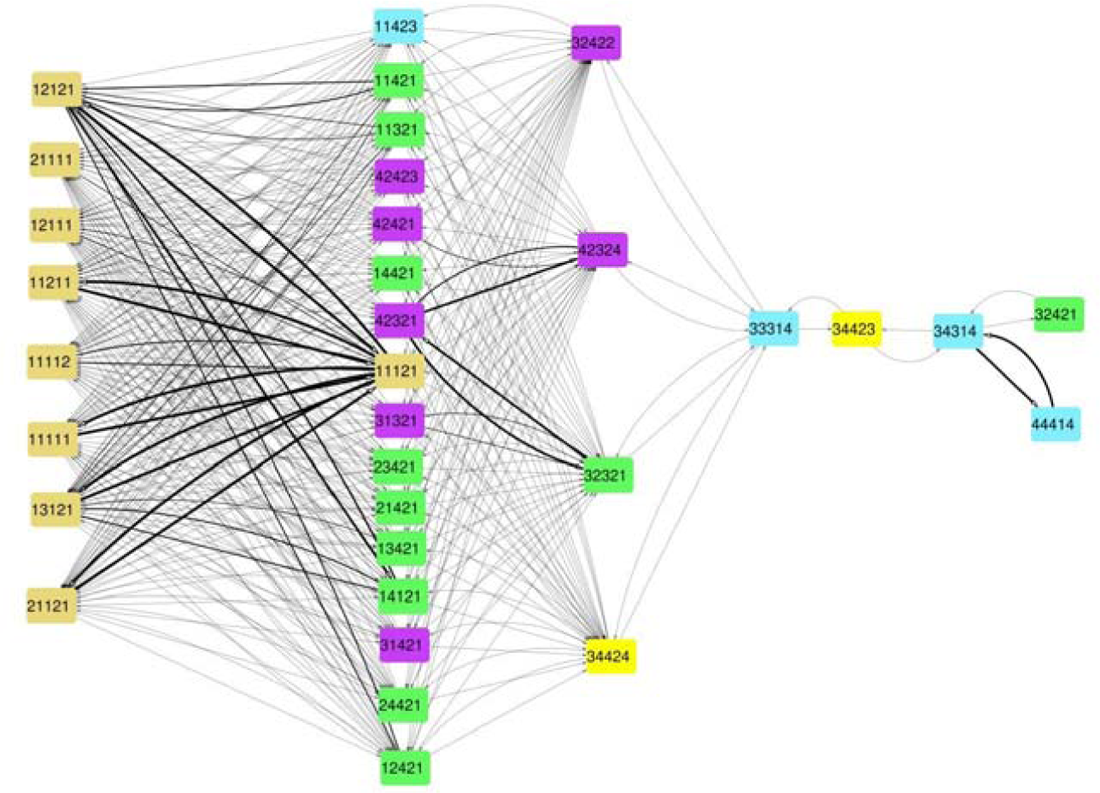
The trend of changes in population in carbon limited scenario. The pink, green, cyan, purple, and yellow circles are individuals with C3-like, C3-C4 intermediate, C2, C4-like, C4 photosynthesis, respectively. Since we drew only individuals with high frequency in the population, this graph is not necessarily connected. Nodes represent the individuals, and the thickness of each arrow represents the “chance” to observe a transition from the first individual to the second one. The width of each edge is proportional to the log frequency transition from the source node to the destination node as explained in Methods section.

### Scenario 4: water limitation

The logarithm of frequency of last generation’s individuals for water limitation scenario is plotted in Fig. 10A, similar to previous Scenarios.

**Figure 10.**
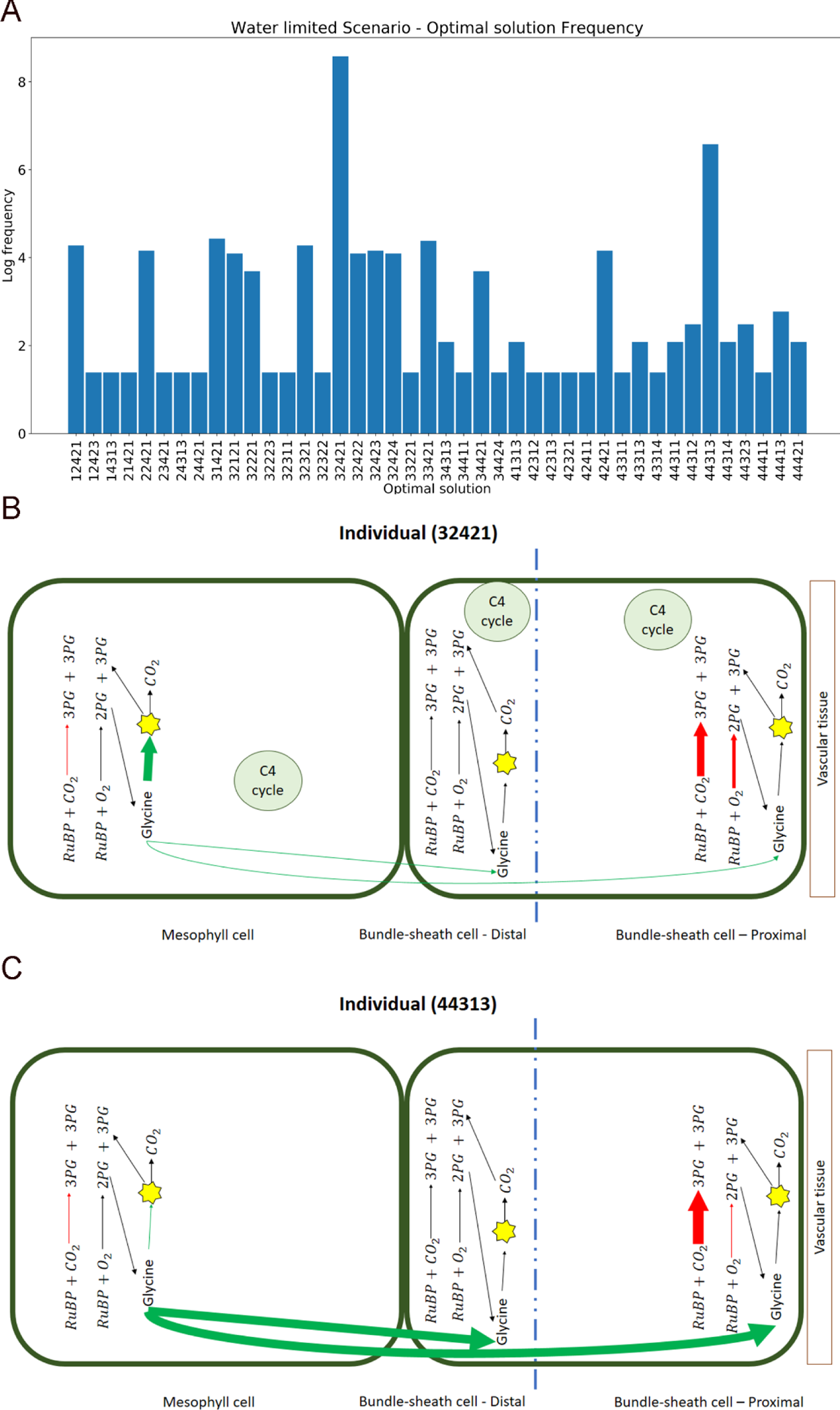
Water limitation scenario. A) The logarithm of frequency of optimal individuals in the last generation; B, C) Schematic representation of traits in simulated metabolic pathways of the top two individuals The thickness of the arrows symbolically indicates the activity of a pathway or reaction in a particular cell. The blue dotted line represents the division of the bundle sheath cell into two proximal and distal parts. The yellow star schematically represents the reactions that convert glycine to 3PG. The size of mesophyll cell and bundle sheath cell as well as the size of proximal and distal parts of bundle sheath cell are shown in proportion to the participation of these parts in the objective function.

The best photosynthetic strategy in the water limitation scenario is C4-like, “32421”, according to Fig. 10B. Except for this individual, the individual named “44313” which is equivalent to a C2 photosynthetic strategy, is the best photosynthetic strategy. We schematically showed the metabolic traits of these two individuals in Fig. 10B and Fig. 10C, respectively.

We showed the evolutionary trajectories of these individuals using a directed graph for water limitation simulation in Fig. 11.

**Figure 11.**
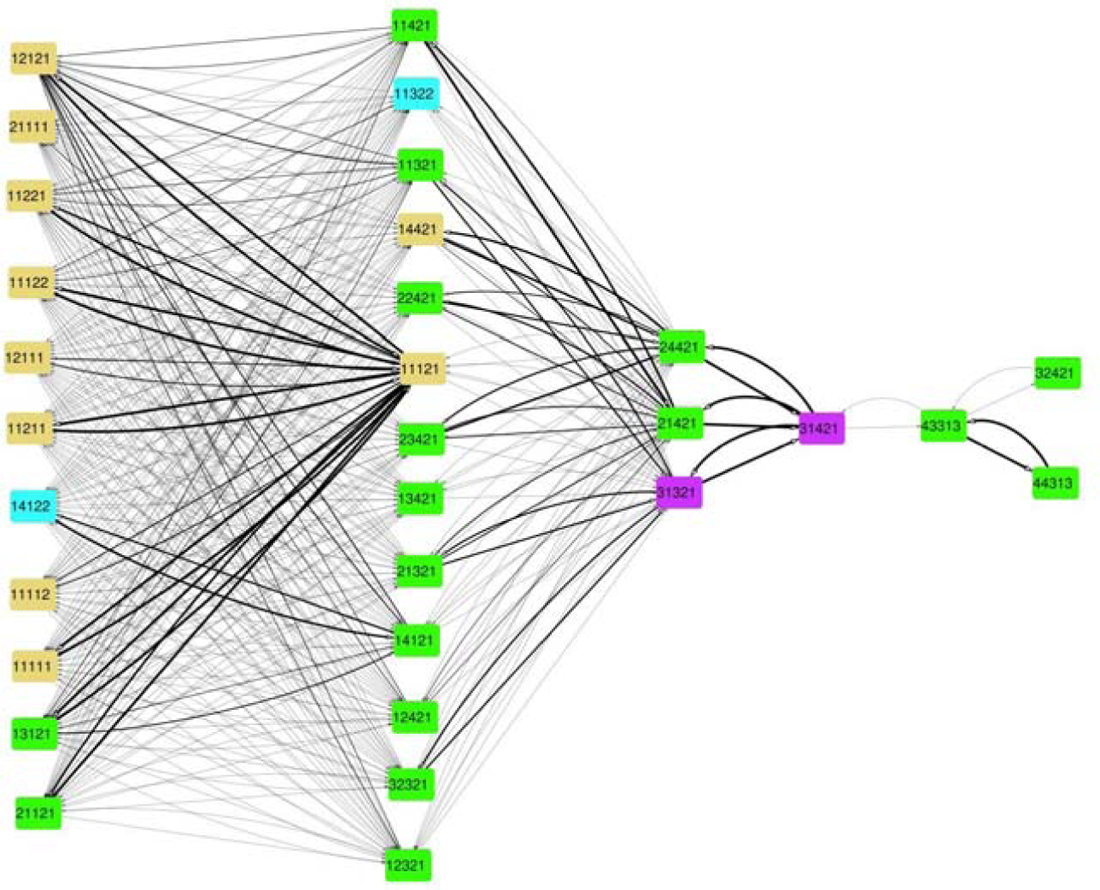
The trend of changes in population in water-limited scenario. The pink, green, cyan, purple, and yellow circles are individuals with C3-like, C3-C4 intermediate, C2, C4-like, C4 photosynthesis, respectively. Since we drew only individuals with high frequency in the population, this graph is not necessarily connected. Nodes represent the individuals, and the thickness of each arrow represents the “chance” to observe a transition from the first individual to the second one. The width of each edge is proportional to the log frequency transition from the source node to the destination node as explained in Methods section.

## DISCUSSION

Several studies have previously utilized constraint-based metabolic network modeling to simulate the outcome of selection pressure on metabolism [17, 20] and the evolutionary trajectories [18]. Recently, CBM is used to predict the optimal photosynthetic metabolic solutions, *i.e.*, metabolic network flux distributions, under different environmental conditions, to investigate the evolution of C4 from C3 photosynthesis [25]. More specifically, the authors developed a two-cell model and investigated the effect of different growth conditions on metabolic fluxes and interpreted flux distributions to find which photosynthetic strategy is used in each condition. They suggested that high photorespiration flux, as well as resource limitation, are selective pressures which drive the C4 evolution. Furthermore, they proposed that the distribution of photons between mesophyll and bundle sheath cells, in addition to the availability of light determines which decarboxylating enzyme in the bundle sheath cells should be active. They suggested that CBM of metabolic networks can be used to study molecular evolution in eukaryotes, and more specifically in plants, with acceptable results.

### Two-cell model captures the essential differences between C3 and C4 photosynthesis

In the present work, we first created a two-cell model based on an existing metabolic network model of *Arabidopsis thaliana*. We should emphasize that instead of examining a single model with a certain parameter setting, as in the work of Blätke and Bräutigam [25], we studied a population of metabolic networks with different P-genotypes using the GA, and then, analyzed the evolutionary trajectories that lead to C4 photosynthesis.

We verified its predictions in C3 and C4 states. Our model successfully predicts that C4 photosynthesis has a higher maximum biomass production capacity in comparison to C3 photosynthesis. Studies such as [12, 26] showed that C4 plants produce more biomass compared to C3 plants. Moreover, it is reported that C4 plants need less Rubisco in their leaves than C3 plants [5, 27]. In line with this observation, our model predicts C4 photosynthesis requires less Rubisco flux than C3 photosynthesis for producing a unit of biomass. Furthermore, it is reported that C4 plants have higher efficiency for conversion of photons into biomass compared to C3 plants [28], and our model predicts that for production of one unit of biomass, the C4 model requires less photons than the C3 model. Additionally, the model predicts that C4 photosynthesis needs less nitrogen for the production of one unit of biomass. It is widely accepted that C4 plants have higher nitrogen use efficiency than C3 plants [29]. Finally, more ATP requirement of C4 pathways is supposed to be a disadvantage of this pathway [30], and our model predicts that C4 photosynthesis needs more ATP to produce one unit of biomass compared to C3 plants. These results showed that our model predictions are (at least qualitatively) in good agreement with the literature.

### Different limitation scenarios may lead to evolution of different photosynthetic features

In the next step, we combined GA, which itself was developed by inspiration from evolution, with CBM to simulate photosynthesis evolution. We chose five photosynthetic-related P-traits with predefined levels, as described in the Methods section. By defining a relevant fitness function, we examined the process of changing these P-traits from the C3 state to more optimal state(s) under different selection scenarios.

Considering changes of each P-trait during the simulations, the similarity between the carbon and water-limited condition was noticeable (Fig. 3 A_iii_-E_iii_ and A_iv_-E_iv_). Interestingly, this observation, which is achieved from metabolic simulations, is consistent with the reported coupling of water and carbon cycles in terrestrial plants [31].

In several studies that have modeled C4 evolution, it is proposed that anatomical enablers like increasing BS number or size are required modifications that assists the evolving C4 trait. It is evident from our simulation study (Fig. 3 C_i_-C_iv_) that the increasing contribution of BS to biomass production in all conditions is favored. We observed an apparent unfavorable presence and activity of Rubisco in the BS in nitrogen-limited conditions (Fig. 3 A_ii_). Our observations are in line with the previous report that the anatomical changes are more affected by nitrogen limitation as leaf’s construction is constrained by nitrogen uptake than water limitation [32].

After the 31st generation (Fig. 3 A_ii_), individuals with the red state (M/BS = 0.1) favored over the green ones (M/BS = 0.01). The observed increased activity of Rubisco in mesophyll is associated with higher contribution of M cells to biomass production (Fig. 3 C_ii_) and evolution of a fully functional glycine pump (Fig. 3E_ii_) in N-limited scenarios compared to C-(Fig. 3 C_iii_, E_iii_) or W-limited scenarios (Fig. 3C_iv_, E_iv_).

Considering the observed pattern for transporting of glycine between the M and the BS cell (Fig. 3 E_i_-E_iv_), one can observe that individuals with a ratio of GDC activity in M to glycine transport to BS of 0.01 were evolved in population. This observation is presumably related to the fact that the limited carbon or nitrogen resources are more associated with the evolution of functional glycine pump and C4 photosynthesis compared to no-limitation condition [33]. It is suggested that the establishment of glycine pump and C2 photosynthesis is an essential initial step in the evolution of C4 photosynthesis [34]. Plants that use such photorespiratory CO_2_ pumps are often referred to as C3-C4 intermediates [13] and are stable evolutionary states toward C4 [35]. Under resource limited conditions, many individuals evolve glycine pumps, in line with this hypothesis. Nevertheless, we noticed that many individuals maintained the C3 state for this P-trait, especially in the water limitation scenario. However, one should bear in mind that the currently accepted hypothesis of C4 evolution is mainly developed based on the observations from a spectrum of C3 and C4 plants in the genus *Flaveria* [34]. Our computational modeling suggests that, in addition to the well-known trajectory of C4 evolution via establishment of glycine pump mechanism, other evolutionary trajectories might also exist that do not necessitate this photorespiratory CO_2_ pump mechanism.

Our observation (Fig. 3D_i_-D_iv_) that activation of C4 metabolic reactions is more favored in nitrogen-limited scenarios than carbon or water scarcity agrees with the findings that less fertile lands drive the evolution of C4 evolution[12].

### Trajectories of C4 photosynthesis evolution in high photorespiration rate conditions

In the next step, we investigated the effects of resource scarcity on the evolutionary outcomes and its impact on evolutionary trajectories of C4 establishment. Our simulations showed that limitation of various resources can lead to different optimal photosynthetic strategies. If carbon or nitrogen are not limited, C4-like photosynthesis predominantly evolves. On the other hand, in an environment in which carbon resource is limited, our model predicts that C2 and C4-like plants have the best performances.

Previous studies have suggested that C2 photosynthesis has evolved in plants and algae at the environment that carbon availability is limited, and the photorespiration rate is high [36]. Furthermore, in an environment in which nitrogen is scarce, our model predicts that C4 and proto-Kranz states are the superior photosynthetic strategies. Our findings are in line with the observation that C4 photosynthesis is a more useful strategy in ecosystems with limitation of nitrogen by having higher productivity and investment in sexual reproduction compared to C3 pathway [12]. Furthermore, we believe that an indirect implication of this finding is that C2 and proto-Kranz states are not just intermediate stages in the evolution of C4 pathway, as they can be the best photosynthetic mechanisms under certain environmental conditions. Recently, it has been suggested that C2 photosynthesis is a stable evolutionary state and should not be considered just as an intermediate stage of C4 evolution [35].

We believe that choosing *A. thaliana* metabolic network model might have influenced the output of our simulations. *A. thaliana* belongs to the *Brassicaceae* family, and according to a study by Schlüter *et al.* [37], although there are many C3-C4 intermediates among *Brassicaceae*, no C4 plant has been discovered in this family yet. In other words, if well-curated CBM of other families that have C4 species was available, we believe the outcome of the simulations might have contained more C4 individuals compared to our simulations.

Many studies suggested that certain genomes have anatomical and metabolic components that facilitate the C4 evolution. For example, in the genus *Flaveria* there are C3, C3-C4 intermediates and C4 species. We speculate that if a well-curated metabolic network of a C3 *Flaveria* species, *e.g., F. cronquistii*, was available, the simulations might have provided better insights into the trajectories toward C4 photosynthesis evolution. To the best of our knowledge, such a metabolic network is not available yet.

## METHODS

### The metabolic model

We used a metabolic network model of *A. thaliana,* which was constructed by Nikoloski et al. [24]. The original model has 549 reactions and 407 metabolites. This model was slightly modified and then utilized as the “base model” in our study. We described our modifications to the original model in S1 Appendix. The two-cell model is constructed by combining two instances of the base model as the two consecutive compartments of the two-cell model, namely mesophyll cell, and bundle sheath cell. All reactions of the bundle sheath cell are duplicated, except for the cytoplasmic and boundary reactions, therefore the bundle sheath compartment of our two-cell model have distal and proximal organelles as shown in Fig 1. The two-cell model is available in the S2 Appendix in mat format.

### FBA

A stoichiometric matrix is a matrix whose rows represent reactions, and its columns represent metabolites. If a metabolic network model has *n* reactions and *m* metabolites, then its stoichiometric matrix is an *n × m* matrix. The dot product of a stoichiometric matrix (*s*) and the vector of reaction fluxes (*v*) equals to the concentration changes in a metabolic network:

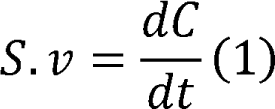

Under steady-state conditions, metabolite concentration changes will be negligible, and therefore, Eq. 1 can be written as:

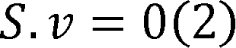

In addition to stoichiometric matrix, other information about the constraints on each reaction can be retrieved from literature or databases. For example, certain reactions may be irreversible, or may have a specific flux limitation. Such “irreversibility” and “capacity” constraints limit the possible range of fluxes through reactions. In other words, each reaction has lower and upper bounds that can be represented as follows:

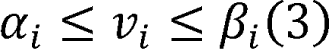

where *α_i_* and *β_i_* are lower and upper bounds, respectively. When lower or upper bounds are not specified, *α_i_* and *β_i_* values are simply assumed to be infinite.

Flux balance analysis (FBA) is a mathematical technique for simulating metabolic fluxes within a metabolic network model under certain environmental conditions [22]. To find a biologically relevant vector of metabolic fluxes, an objective function is defined, which represents the metabolic goal of the cell to be optimized. Maximizing the biomass production rate is widely used in FBA studies [9, 38]. In FBA, linear programming is used to solve an optimization problem to find a vector of feasible fluxes (ν). The optimization problem can be written as follows:

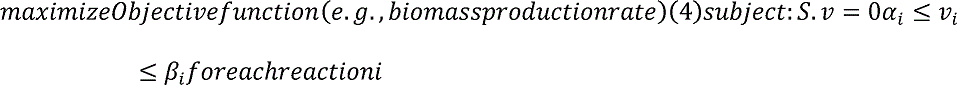

The COnstraint-Based Reconstruction and Analysis (COBRA) Toolbox is a set of functions that can be used to reconstruct and analyze metabolic network model [39]. We used COBRA Toolbox to perform our FBA analysis.

As mentioned above, we used a previously published metabolic network of *A. thaliana* [24] as our base model. We set constraints based on the previous studies on the reaction fluxes [23, 24]. The objective function of the two-cell model was defined as Equation 5.

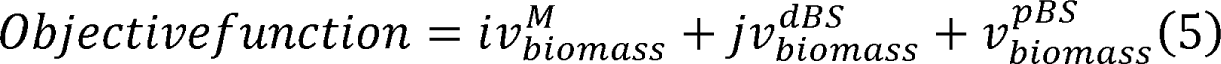

In Equation 5, *ν^M^_biomass_*, *ν^dBS^_biomass_*, and *ν^pBS^_biomass_* represent the biomass reaction of mesophyll, distal- and proximal-bundle sheath, respectively. Here, *i* and *j* are fixed coefficients which define the contribution of mesophyll, distal- and proximal bundle sheath to produce the leaf biomass, respectively. We set *i* and *j* according to the third trait of each individual to represent a specific photosynthetic strategy.

### Genetic algorithm (GA) design

We used a GA to see how the two-cell models evolve under different environmental conditions. To define the GA problem, first, we should decide a set of *n* “P-traits”, *i.e.*, photosynthetic traits, to be optimized in our simulations and specify the acceptable levels for each of them. A P-genotype can be defined as an ordered *n*-tuple of the P-traits. Each P-genotype has a biological P-phenotype. Additionally, for each P-genotype, one can consider a corresponding metabolic model, based on which a fitness value can be computed for that certain individual.

Based on previous studies, five P-traits were chosen [7, 8, 14, 40]. These P-traits are selected in a way that they can capture the main differences of C3 and C4 photosynthetic strategies. Using literature, we defined levels for these traits to represent ‘C3’, ‘C3-C4 intermediate’, and ‘C4’ states. These levels and the photosynthetic states, which they represent are summarized in Table 2.

A GA consists of three steps: (i) creating the initial population; (ii) selection of the best individuals based on a fitness function; and (iii) creating the individuals of the next generation using operations such as mutation, crossover, or elitism. We created a population of C3 and C3-like individuals for our initial population. For each P-genotype, a corresponding metabolic network was constructed, and the optimal value of the objective function was computed as a proxy for its fitness value.

The fitness of each individual was calculated from its objective function (a linear combination of biomass productions of mesophyll and bundle sheath cell), ATP consumption, and sum of all metabolic fluxes:

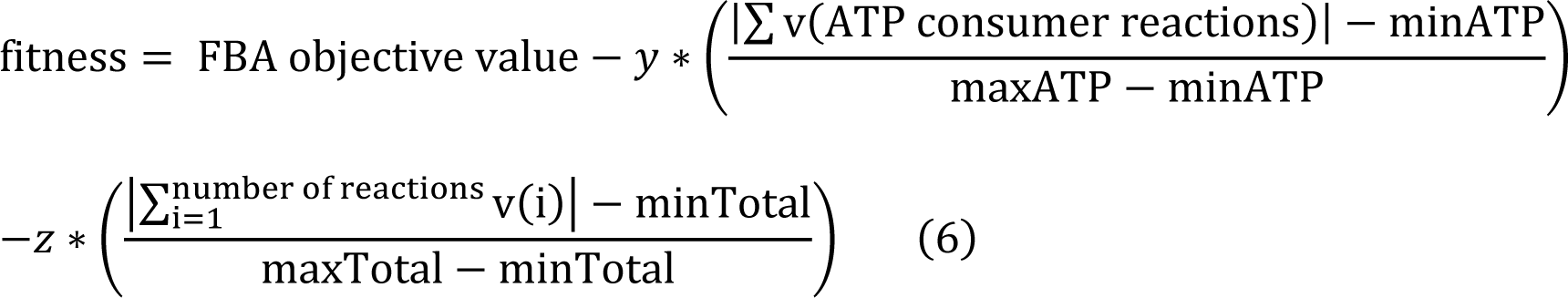

Each individual, based on its fitness score, has a chance to create the individuals of next generation. We have discussed, in details, the steps of implementing the GA in the S1 Appendix.

### Simulation of evolutionary events

To predict the effects of resource scarcity on the evolutionary trajectory and outcome of simulation, we designed four scenarios for simulation. In the first scenario, there is no limitation on resource availability for the models. In the next scenarios, we limited the carbon, nitrogen, and water availability to the models, respectively. We emphasize that in each of these scenarios, only one resource (namely water, nitrogen, and carbon) was limited. The detailed procedure is mentioned in the S1 Appendix. The codes that were used for these simulations are available in S3 Appendix.

### Analyzing the pattern of changes in individual traits under each scenario

To find patterns in the transition of genotypes during the evolution for each scenario, we count the occurrence of each genotype in each generation for a specific scenario in all repetitions of that scenario. Then, we create a directed graph using the Cytoscape [41] which its nodes represent the genotypes and its edges’ width was proportional to the log frequency of transition from the origin node to the destination node. Moreover, we clustered the genotypes from first to the tenth generation into one group and show them as a single column. We repeated this for the next nine intervals. To calculate the transition matrix, we saved the parents’ information for each individual when the GA executed and counted the number of transitions from each parent to each child in all generations of a specific scenario in all repetitions and stored its value in a matrix. Frequency of each individual in each generation of our simulations and the transition matrix are available in S4 Appendix.

## ACKNOWLEDGMENTS

We would like to thank Ilia Abolhasani (Sharif University of Technology) for his help with the implementation of the genetic algorithm. We would also like to thank Hamideh Fouladiha, Shima GhaediZadeh, Mohmmad-Hossein Motealehi-Ardakani, Sevan Gholipour, and Meisam Yousefi (University of Tehran) for their helpful discussions and comments on this research project. A.M.B-M gratefully acknowledges the financial support from the Iran National Science Foundation (INSF) [funding reference number 95838484].

## SUPPORTING INFORMATION CAPTIONS

S1 Appendix. Step by step explanation of the simulation procedures

S2 Appendix. Our two-cell model in the`.mat` format

S3 Appendix. The source codes used for simulations in the present work.

S4 Appendix. Tables of each individual’s frequency in each generation of simulations and transition matrix

## REFERENCES

1. South PF, Cavanagh AP, Liu HW, Ort DR. Synthetic glycolate metabolism pathways stimulate crop growth and productivity in the field. Science (New York, NY). 2019;363(6422):eaat9077. Epub 2019/01/05. doi: 10.1126/science.aat9077. PubMed PMID: 30606819; PubMed Central PMCID: PMCPMC7745124.

2. Cui H. Challenges and Approaches to Crop Improvement Through C3-to-C4 Engineering. Frontiers in plant science. 2021;12. doi: 10.3389/fpls.2021.715391.

3. Flügel F, Timm S, Arrivault S, Florian A, Stitt M, Fernie AR, et al. The Photorespiratory Metabolite 2-Phosphoglycolate Regulates Photosynthesis and Starch Accumulation in Arabidopsis. The Plant Cell. 2017;29(10):2537–51. doi: 10.1105/tpc.17.00256.

4. Zhu X-G, Long SP, Ort DR. What is the maximum efficiency with which photosynthesis can convert solar energy into biomass? Current Opinion in Biotechnology. 2008;19(2):153–9. doi: https://doi.org/10.1016/j.copbio.2008.02.004.

5. Schlüter U, Weber AP. The Road to C4 Photosynthesis: Evolution of a Complex Trait via Intermediary States. Plant & cell physiology. 2016;57(5):881–9. Epub 2016/02/20. doi: 10.1093/pcp/pcw009. PubMed PMID: 26893471.

6. Fouracre JP, Ando S, Langdale JA. Cracking the Kranz enigma with systems biology. Journal of Experimental Botany. 2014;65(13):3327–39. doi: 10.1093/jxb/eru015%J Journal of Experimental Botany.

7. Sage RF, Khoshravesh R, Sage TL. From proto-Kranz to C4 Kranz: building the bridge to C4 photosynthesis. Journal of Experimental Botany. 2014;65(13):3341–56. doi: 10.1093/jxb/eru180.

8. Bräutigam A, Gowik U. Photorespiration connects C3 and C4 photosynthesis. Journal of Experimental Botany. 2016;67(10):2953–62. doi: 10.1093/jxb/erw056.

9. Mallmann J, Heckmann D, Bräutigam A, Lercher MJ, Weber APM, Westhoff P, et al. The role of photorespiration during the evolution of C4 photosynthesis in the genus Flaveria. eLife. 2014;3:e02478. doi: 10.7554/eLife.02478.

10. Heckmann D. C4 photosynthesis evolution: the conditional Mt. Fuji. Curr Opin Plant Biol. 2016;31:149–54. Epub 2016/05/07. doi: 10.1016/j.pbi.2016.04.008. PubMed PMID: 27153468.

11. Sage TL, Busch FA, Johnson DC, Friesen PC, Stinson CR, Stata M, et al. Initial Events during the Evolution of C4 Photosynthesis in C3 Species of Flaveria. Plant Physiology. 2013;163(3):1266–76. doi: 10.1104/pp.113.221119.

12. Ripley BS, Abraham TI, Osborne CP. Consequences of C4 photosynthesis for the partitioning of growth: a test using C3 and C4 subspecies of Alloteropsis semialata under nitrogen-limitation. Journal of Experimental Botany. 2008;59(7):1705–14. doi: 10.1093/jxb/erm210.

13. Stata M, Sage TL, Sage RF. Mind the gap: the evolutionary engagement of the C4 metabolic cycle in support of net carbon assimilation. Current Opinion in Plant Biology. 2019;49:27–34. doi: https://doi.org/10.1016/j.pbi.2019.04.008.

14. Edwards EJ. Evolutionary trajectories, accessibility and other metaphors: the case of C4 and CAM photosynthesis. New Phytologist. 2019;223(4):1742–55. doi: https://doi.org/10.1111/nph.15851.

15. Schuler ML, Mantegazza O, Weber AP. Engineering C4 photosynthesis into C3 chassis in the synthetic biology age. The Plant journal: for cell and molecular biology. 2016;87(1):51–65. Epub 2016/03/08. doi: 10.1111/tpj.13155. PubMed PMID: 26945781.

16. Schlüter U, Weber APM. Regulation and Evolution of C(4) Photosynthesis. Annual review of plant biology. 2020;71:183–215. Epub 2020/03/07. doi: 10.1146/annurev-arplant-042916-040915. PubMed PMID: 32131603.

17. Pál C, Papp B, Lercher MJ, Csermely P, Oliver SG, Hurst LD. Chance and necessity in the evolution of minimal metabolic networks. Nature. 2006;440(7084):667-70. doi: 10.1038/nature04568.

18. Yizhak K, Tuller T, Papp B, Ruppin E. Metabolic modeling of endosymbiont genome reduction on a temporal scale. Molecular Systems Biology. 2011;7(1):479. doi: https://doi.org/10.1038/msb.2011.11.

19. Schönknecht G, Weber AP, Lercher MJ. Horizontal gene acquisitions by eukaryotes as drivers of adaptive evolution. BioEssays: news and reviews in molecular, cellular and developmental biology. 2014;36(1):9–20. Epub 2013/12/11. doi: 10.1002/bies.201300095. PubMed PMID: 24323918.

20. Pál C, Papp B, Lercher MJ. Adaptive evolution of bacterial metabolic networks by horizontal gene transfer. Nature Genetics. 2005;37(12):1372–5. doi: 10.1038/ng1686.

21. Szappanos B, Fritzemeier J, Csörgő B, Lázár V, Lu X, Fekete G, et al. Adaptive evolution of complex innovations through stepwise metabolic niche expansion. Nature Communications. 2016;7(1):11607. doi: 10.1038/ncomms11607.

22. Orth JD, Thiele I, Palsson BØ. What is flux balance analysis? Nature Biotechnology. 2010;28(3):245–8. doi: 10.1038/nbt.1614.

23. Robaina-Estévez S, Daloso DM, Zhang Y, Fernie AR, Nikoloski Z. Resolving the central metabolism of Arabidopsis guard cells. Scientific Reports. 2017;7(1):8307. doi: 10.1038/s41598-017-07132-9.

24. Arnold A, Nikoloski Z. Bottom-up Metabolic Reconstruction of Arabidopsis and Its Application to Determining the Metabolic Costs of Enzyme Production. Plant Physiology. 2014;165(3):1380–91. doi: 10.1104/pp.114.235358.

25. Blätke M-A, Bräutigam A. Evolution of C4 photosynthesis predicted by constraint-based modelling. eLife. 2019;8:e49305. doi: 10.7554/eLife.49305.

26. Rubio G, Gutierrez Boem FH, Lavado RS. Responses of C3 and C4 grasses to application of nitrogen and phosphorus fertilizer at two dates in the spring. Grass and Forage Science. 2010;65(1):102–9. doi: https://doi.org/10.1111/j.1365-2494.2009.00728.x.

27. Reyna-Llorens I, Hibberd JM. Recruitment of pre-existing networks during the evolution of C4 photosynthesis. Philosophical transactions of the Royal Society of London Series B, Biological sciences. 2017;372(1730):20160386. doi: doi:10.1098/rstb.2016.0386.

28. Wang C, Guo L, Li Y, Wang Z. Systematic Comparison of C3 and C4 Plants Based on Metabolic Network Analysis. BMC Systems Biology. 2012;6(2):S9. doi: 10.1186/1752-0509-6-S2-S9.

29. Sage RF. A portrait of the C4 photosynthetic family on the 50th anniversary of its discovery: species number, evolutionary lineages, and Hall of Fame. Journal of Experimental Botany. 2016;67(14):4039–56. doi: 10.1093/jxb/erw156.

30. Gowik U, Westhoff P. The path from C3 to C4 photosynthesis. Plant Physiol. 2011;155(1):56–63. Epub 2010/10/14. doi: 10.1104/pp.110.165308. PubMed PMID: 20940348; PubMed Central PMCID: PMCPMC3075750.

31. Gentine P, Green JK, Guérin M, Humphrey V, Seneviratne SI, Zhang Y, et al. Coupling between the terrestrial carbon and water cycles—a review. Environmental Research Letters. 2019;14(8):083003. doi: 10.1088/1748-9326/ab22d6.

32. Farrior CE, Tilman D, Dybzinski R, Reich PB, Levin SA, Pacala SW. Resource limitation in a competitive context determines complex plant responses to experimental resource additions. Ecology. 2013;94(11):2505–17. doi: https://doi.org/10.1890/12-1548.1.

33. Ripley BS, Cunniff J, Osborne CP. Photosynthetic acclimation and resource use by the C3 and C4 subspecies of Alloteropsis semialata in low CO2 atmospheres. 2013;19(3):900-10. doi: https://doi.org/10.1111/gcb.12091.

34. Monson RK, Edwards GE, Ku MSB. C3-C4 Intermediate Photosynthesis in Plants. BioScience. 1984;34(9):563–74. doi: 10.2307/1309599.

35. Lundgren MR. C2 photosynthesis: a promising route towards crop improvement? New Phytologist. 2020;228(6):1734–40. doi: https://doi.org/10.1111/nph.16494.

36. Hagemann M, Fernie AR, Espie GS, Kern R, Eisenhut M, Reumann S, et al. Evolution of the biochemistry of the photorespiratory C2 cycle. Plant biology (Stuttgart, Germany). 2013;15(4):639–47. Epub 2012/12/04. doi: 10.1111/j.1438-8677.2012.00677.x. PubMed PMID: 23198988.

37. Schlüter U, Bräutigam A, Gowik U, Melzer M, Christin P-A, Kurz S, et al. Photosynthesis in C3–C4 intermediate Moricandia species. Journal of Experimental Botany. 2016;68(2):191–206. doi: 10.1093/jxb/erw391.

38. Schulze S, Westhoff P, Gowik U. Glycine decarboxylase in C3, C4 and C3-C4 intermediate species. Curr Opin Plant Biol. 2016;31:29-35. Epub 2016/04/03. doi: 10.1016/j.pbi.2016.03.011. PubMed PMID: 27038285.

39. Heirendt L, Arreckx S, Pfau T, Mendoza SN, Richelle A, Heinken A, et al. Creation and analysis of biochemical constraint-based models using the COBRA Toolbox v.3.0. Nature Protocols. 2019;14(3):639-702. doi: 10.1038/s41596-018-0098-2.

40. Sage RF, Monson RK, Ehleringer JR, Adachi S, Pearcy RW. Some like it hot: the physiological ecology of C4 plant evolution. Oecologia. 2018;187(4):941–66. doi: 10.1007/s00442-018-4191-6.

41. Shannon P, Markiel A, Ozier O, Baliga NS, Wang JT, Ramage D, et al. Cytoscape: a software environment for integrated models of biomolecular interaction networks. Genome research. 2003;13(11):2498–504. Epub 2003/11/05. doi: 10.1101/gr.1239303. PubMed PMID: 14597658; PubMed Central PMCID: PMCPMC403769.

