## Supplementary material for "Computational modeling of the evolutionary transition from C3 to C4 photosynthesis": S1 Appendix

**Short Title: Modeling the evolutionary transition from C3 to C4 photosynthesis**

Armin Dadras^1^, Sayed-Amir Marashi^1,*^, Kaveh Kavousi^2^, Ali-Mohammad Banaei-Moghaddam^3,*^

^1^ Department of Biotechnology, College of Science, University of Tehran, Tehran, Iran

^2^ Laboratory of Complex Biological Systems and Bioinformatics (CBB), Institute of Biochemistry and Biophysics (IBB), University of Tehran, Tehran, Iran

^3^ Laboratory of Genomics and Epigenomics (LGE), Department of Biochemistry, Institute of Biochemistry and Biophysics (IBB), University of Tehran

* Corresponding authors

### (S.A.M.), (A.M.B-M)

### S1 Appendix: Step by step explanation of the simulation

First, we removed unrealistic reactions that directly connect organelles to the outside of the cell. Since compartmentalization of the present model is important, such reactions could have resulted in wrong predictions.

In the next step, a list of C4-related reactions was compiled, according to the literature [1-4], Then, we checked whether all these C4-related reactions are present in our model, to add the missing reactions to the model. A list of reactions that are added to the model are presented in Table 1.

**Table 1. List of added C4-related reactions to the model**

| **Reaction name** | **Reaction formula** | **Reference** |
| --- | --- | --- |
| Malate-Oxaloacetate shuttle II | OAA[c]+Mal[h] -> OAA[h] + Mal[c] | [5] |
| Alanine transaminase | KG[c] + Ala[c] <=> Pyr[c] + Glu[c] | [6] |
| Aspartate Aminotransferase, mitochondrial | KG[m] + Asp[m] <=> OAA[m] + Glu[m] | [7] |
| Mal/KG DTC | KG[m] + Mal[c] -> KG[c] +Mal[m] | [8] |
| Asp-Glu AGCs | Glu[c] + Asp[m] <=> Glu[m] + Asp[c] | [8] |
| NAD-dependent malic enzyme II, mitochondrial | Mal[m] + NAD[m] -> CO2[m] + NADH[m] + Pyr[m] | [9] |

Moreover, according to [10, 11], we added certain constraints on reaction fluxes in Table 2.

**Table 2. List of the constraints that were set on the model**

| **Reaction name** | **Lower bound** | **Upper bound** | **Reaction name** | **Lower bound** | **Upper bound** |
| --- | --- | --- | --- | --- | --- |
| G6PDH_h | 0 | 0 | Ex_Asn_c | 0 | 0 |
| PPIF6PK_c | 0 | 0 | Ex_Asp_c | 0 | 0 |
| MalEx | 0 | 0 | Ex_Cys_c | 0 | 0 |
| Ex_Ala_c | 0 | 0 | Ex_Gln_c | 0 | 0 |
| Ex_Arg_c | 0 | 0 | Ex_Glu_c | 0 | 0 |
| Ex_Leu_c | 0 | 0 | Ex_Gly_c | 0 | 0 |
| Ex_Lys_c | 0 | 0 | Ex_His_c | 0 | 0 |
| Ex_Met_c | 0 | 0 | Ex_Ile_c | 0 | 0 |
| Ex_Phe_c | 0 | 0 | Ex_Val_c | 0 | 0 |
| Ex_Pro_c | 0 | 0 | Ex_Tre_c | 0 | 0 |
| Ex_Ser_c | 0 | 0 | Ex_Thr_c | 0 | 0 |
| Ex_Trp_c | 0 | 0 | Ex_Tyr_c | 0 | 0 |
| Ex_glc | 0 | 0 | Ex_cellulose | 0 | 0 |
| Ex_Frc | 0 | 0 | Ex_Mas | 0 | 0 |
| Ex_Suc | 0 | 0 | Ex_MACP | 0 | 0 |
| Bio_CLim | 0 | 0 | Bio_NLim | 0 | 0 |
| Bio_AA | 0 | 0 | NGAM_c | 0.001 | 1000 |
| NGAM_h | 0.001 | 1000 | NGAM_m | 0.001 | 1000 |
| FBPase_h | 0.001 | 1000 | SBPA_h | 0.001 | 1000 |

For the carboxylase and oxygenase activities of Rubisco, there are two separate reactions in the model [10], which means that these two reactions are practically uncoupled in the model. In order to couple carboxylase and oxygenase fluxes and to have a fixed ratio of carboxylase to oxygenase activities, we removed these two reactions and added a new reaction for the Rubisco to the model, as suggested in [12]:

$$r CO_{2} + r H_{2}O + \left( r + 1 \right)\mathrm{RuBP} +O_{2} \to\left( 2r + 1 \right)\mathrm{PGA} + 2PG (1)$$

The ratio, $r$, can be estimated from CO_2_ pressure in the atmosphere. Liu et al. [12] have calculated this parameter for eight different atmospheric CO_2_ concentrations for *Arabidopsis thaliana*.

**Table 3. The ratio of carboxylation to oxygenation reaction of Rubisco under various CO_2_ concentrations**

| **CO_2_ concentration** | **r** |
| --- | --- |
| 100 ppm | 1.139 |
| 380 ppm | 4.33 |
| 500 ppm | 5.69 |
| 600 ppm | 6.83 |
| 700 ppm | 7.97 |
| 800 ppm | 9.11 |
| 900 ppm | 10.25 |
| 1000 ppm | 11.39 |

In the next step, we created a two-cell model which includes two instances of the modified metabolic network. Many studies such as [13] have reported that proximal and distal organelles in the bundle sheath cells of C4 plants have different characteristics. In order to create a bundle sheath model which has distal and proximal organelles, we divided reactions into two groups.

To represent environmental conditions that increase the photorespiration rate, we set a ratio of carboxylase to oxygenase activity of Rubisco in mesophyll and distal bundle sheath to 1.139 based on experimental data [12]. This ratio for Rubisco in proximal bundle sheath was defined according to each model's parameters. Next, we merged the mesophyll and bundle sheath cell models into a two-cell model to serve as the framework for subsequent steps. Then, we added reactions that transport metabolites between mesophyll and bundle sheath models, according to [2, 3, 14]. The list of reactions that were added to the two-cell model and their formula are presented in Table 4.

**Table 4. List of reactions that transport metabolites between two cells**

| **Reaction name** | **Reaction formula** |
| --- | --- |
| Water transport | M_H2O[c] <=> BS_H2O[c] |
| Oxygen transport | M_O2[c] <=> BS_O2[c] |
| Pi transport | M_Pi[c] <=> BS_Pi[c] |
| SO_4_ transport | M_SO4[c] <=> BS_SO4[c] |
| H_2_S transport | M_H2S[c] <=> BS_H2S[c] |
| Glycolate transport | M_GCA[c] <=> BS_GCA[c] |
| Glyceraldehyde-3-phosphate Transport | M_GAP[c] <=> BS_GAP[c] |
| Malate Transport | M_Mal[c] <=> BS_Mal[c] |
| Pyruvate Transport | M_Pyr[c] <=> BS_Pyr[c] |
| PEP Transport | M_PEP[c] <=> BS_PEP[c] |
| 2-phosphoglycerate Transport | M_2PGA[c] <=> BS_2PGA[c] |
| 3-phosphoglycerate Transport | M_PGA[c] <=> BS_PGA[c] |
| Dihydroxyacetone phosphate Transport | M_DHAP[c] <=> BS_DHAP[c] |
| Sucrose Transport | M_Suc[c] <=> BS_Suc[c] |
| Fructose Transport | M_Frc[c] <=> BS_Frc[c] |
| Glucose Transport | M_Glc[c] <=> BS_Glc[c] |
| Alpha-ketoglutarate Transport | M_KG[c] <=> BS_KG[c] |
| Arginine Transport | M_Arg[c] <=> BS_Arg[c] |
| Histidine Transport | M_His[c] <=> BS_His[c] |
| Lysine Transport | M_Lys[c] <=> BS_Lys[c] |
| Aspartate Transport | M_Asp[c] <=> BS_Asp[c] |
| Glutamate Transport | M_Glu[c] <=> BS_Glu[c] |
| Serine Transport | M_Ser[c] <=> BS_Ser[c] |
| Threonine Transport | M_Thr[c] <=> BS_Thr[c] |
| Asparagine Transport | M_Asn[c] <=> BS_Asn[c] |
| Glutamine Transport | M_Gln[c] <=> BS_Gln[c] |
| Cysteine Transport | M_Cys[c] <=> BS_Cys[c] |
| Glycine Transport | M_Gly[c] <=> BS_Gly[c] |
| Proline Transport | M_Pro[c] <=> BS_Pro[c] |
| Alanine Transport | M_Ala[c] <=> BS_Ala[c] |
| Valine Transport | M_Val[c] <=> BS_Val[c] |
| Isoleucine Transport | M_Ile[c] <=> BS_Ile[c] |
| Leucine Transport | M_Leu[c] <=> BS_Leu[c] |
| Methionine Transport | M_Met[c] <=> BS_Met[c] |
| Phenylalanine Transport | M_Phe[c] <=> BS_Phe[c] |
| Tyrosine Transport | M_Tyr[c] <=> BS_Tyr[c] |
| Tryptophan Transport | M_Trp[c] <=> BS_Trp[c] |

In the next step, we modified boundary reactions to the two-cell model. Inorganic metabolites were only allowed to enter into the bundle sheath cell from outside of the model to simulate the transport of inorganic substances from xylem to the bundle sheath cells in the leaves. Then, these metabolites can be transported from this cell to the mesophyll cell. Boundary reactions that are added to the model, their formula and their lower and upper bounds are shown in Table 5.

**Table 5. List of boundary reactions that were added to the model and their constraints**

| **Reaction name** | **Reaction formula** | **Lower bound** | **Upper bound** |
| --- | --- | --- | --- |
| M Photon input | -> M_hnu[h] | 0 | 1000 |
| BS Photon input | -> BS_hnu[h] | 0 | 1000 |
| D Photon input | -> P_hnu[h] | 0 | 1000 |
| M CO_2_ input | <=> M_CO2[c] | -1000 | 1000 |
| BS CO_2_ input | <=> BS_CO2[c] | -1000 | 1000 |
| BS H_2_O input | <=> BS_H2O[c] | -1000 | 1000 |
| BS Pi input | 3 BS_ATP[c] + 3 BS_H2O[c] ->  2 BS_H[h] + BS_Pi[h] + 3 BS_ADP[c] + 3  BS_Pi[c] | 0 | 1000 |
| D Pi input | 3 BS_ATP[c] + 3 BS_H2O[c] ->  2 P_H[h] + P_Pi[h] + 3 BS_ADP[c] + 3  BS_Pi[c] | 0 | 1000 |
| BS NO_3_ input | 2 BS_ATP[c] + 2 BS_H2O[c] ->  2 BS_ADP[c] + 2 BS_H[c] + 2 BS_Pi[c] +  BS_NO3[c] | 0 | 1000 |
| BS NH_4_ input | -> BS_NH4[c] | 0 | 1000 |
| BS SO_4_ input | 3 BS_ATP[c] + 3 BS_H2O[c] -> 3 BS_ADP[c]  + 3 BS_H[c] + 3 BS_Pi[c] + BS_SO4[c] | 0 | 1000 |
| BS H_2_S input | -> BS_H2S[c] | 0 | 1000 |
| M O_2_ exchange | M_O2[c] <=> | -1000 | 1000 |
| BS O_2_ exchange | BS_O2[c] <=> | -1000 | 1000 |

Suppose that 10% of CO_2_ is lost due to the diffusion, as shown in Equation 2.

$${CO_{2}}_{internal}\to{0.9 CO_{2}}_{internal}+{0.1 CO_{2}}_{external} \left( 2 \right)$$

In order to enable simulations of CO_2_ loss, we first create a list of reactions that produce CO_2_ in our model. For each such reaction, we updated the model as follows. Let Equation 3 represent a hypothetical CO_2_-producing reaction.

$$A\to B+{CO_{2}}_{internal} \left( 3 \right)$$

To consider the effect of CO_2_ loss, we modified every CO_2_-producing reaction in mesophyll, distal bundle sheath, and cytoplasmic bundle sheath according to Equation 4.

$$A\to B+{0.9 CO_{2}}_{internal}+{0.1 CO_{2}}_{external} (4)$$

The objective function of the final two-cell model was defined as Equation 5.

$$Objective function = i v_{\mathrm{biomass}}^{M} + j v_{\mathrm{biomass}}^{\mathrm{dBS}} + v_{\mathrm{biomass}}^{\mathrm{pBS}} (5)$$

In equation 5,$v_{\mathrm{biomass}}^{M}$, $v_{\mathrm{biomass}}^{\mathrm{dBS}}$, and $v_{\mathrm{biomass}}^{\mathrm{pBS}}$ represent the biomass reaction of mesophyll, distal- and proximal-bundle sheath, respectively. Also, *i* and *j* define the contribution of mesophyll, distal- and proximal bundle sheath to produce the leaf biomass, respectively. We set *i* and *j* according to Table 6, trait 3, to represent a specific photosynthetic strategy.

#### Genetic Algorithm (GA) design

To define the GA problem, first, we should decide which traits to be optimized in our simulations and specify the acceptable levels for each of them. According to previous studies and metabolic network limitations, we chose five traits [13, 15-17]. Using literature, we define levels for these traits to represent C3, C3-C4 intermediate and C4 state. These levels and the photosynthetic state, which they represent is shown in Table 6.

**Table 6. Levels of traits in the GA and their photosynthetic equivalents**

| Trait 1: The ratio of Rubisco reaction’s flux in mesophyll cell to bundle sheath cell | | | | |
| --- | --- | --- | --- | --- |
| Trait values | 100 | 10 | 0.1 | 0.01 |
| Considered as | C3 | C3-C4 | C4 | C4 |
| Trait 2: The ratio of carboxylase to oxygenase activity of the Rubisco reaction in proximal chloroplast of bundle sheath cell | | | | |
| Trait values | 1.139 | 1.937 | 3.53 | 4.13 |
| Considered as | C3 | C3-C4 | C3-C4 | C4 |
| Trait 3: Contribution of the mesophyll, distal- and proximal bundle sheath in the production of leaf biomass | | | | |
| Trait values | 8.6, 100 | 6.55, 10 | 2.85, 0.1 | 1, 0.01 |
| Considered as | C3 | C3-C4 | C4 | C4 |
| Trait 4: Upper bound of selected C4 reactions | | | | |
| Trait values | 0 | | 1000 | |
| Considered as | C3 | | C4 | |
| Trait 5: Ratio of “Glycine Decarboxylase” reaction’s flux in the mesophyll cell to “Glycine Transport” reaction’s flux (from the mesophyll cell to the bundle sheath cell) | | | | |
| Trait values | 100 | 10 | 0.1 | 0.01 |
| Considered as | C3 | C3-C4 | C3-C4 | C4 |

For the first and fifth traits, we consider four ratios which represent enormously greater, greater, lesser, and extremely lesser. For the second trait, we supposed that the relation between r and CO_2_ concentration is linear and we chose two CO_2_ concentrations from [12] to represent our lowest and highest levels and interpolated two regularly spaced points between these extreme to symbolize C3-C4 intermediates. For the third trait, we used the fact that the average ratio of mesophyll to bundle sheath cells area in cross-section of leaves is 8.6 for C3 and in the range of 1 to 4 for C4 plants [18]. We Considered 8.6 and 1 as our two extremes and chose two regularly spaced points between these two extremes. It is important to note that the third level of this trait is in the C4 range. We supposed that our fourth trait is binary. We assumed that fluxes of the selected reactions from C4 pathway are negligible in C3 plants and therefore can be considered as zero compared to the C4 plants [2, 19, 20]. It is important to emphasize that considering different traits or different levels for these traits could change the result of the simulation.

Due to the nature of the GA, if two chromosomes have the same fitness value which is high, but one of them appeared in the earlier generations, then this chromosome has more contribution to the final population make up. To eliminate this bias, we performed a hundred replications for each scenario. It should be noted that in all scenarios we assumed that the photorespiration rate is high regardless of resource availability.

Based on our literature review and model reactions, we chose reactions listed in Table 7 and changed their upper bound according to the fourth trait’s level [1, 21, 22].

**Table 7. List of putative C4 reactions whose fluxes are constrained**

| **Reaction name in the model** | **Reaction name in the model** |
| --- | --- |
| PEPC1_c | MalDH2NADP_c |
| PEPC2_c | MalDH2NADP_h |
| PyrPiDK_h | M_Malate Oxaloacetate shuttle II_h |
| AlaTA_h | M_Aspartate aminotransferase_m |
| AlaTA_p | M_Malate KG DTC_m |
| AlaTA_m | M_Asp Glu AGCs_m |
| Alanine transaminase | M_NAD-ME II mitochondrial_m |

Because we aimed to simulate the evolution of C4 from C3 photosynthesis, we could not create an initial population with high diversity for the GA. On the other hand, if all individual has the same traits’ level, we would not see any difference in the fitness of individuals which results in defect for the selection step of the GA. Therefore, we create an initial population that all of the individuals had C3 levels for every trait. Then, for 90% of this population, we select one trait randomly and change its level to the second level.

To calculate the fitness of each chromosome, we considered three parameters which are the objective function (equation 5), the sum of the absolute value of fluxes for reactions that consume ATP, and the sum of the absolute value of fluxes for all reactions. Equation 6 shows how we combine these parameters to calculate fitness.

$$fitness= Objective function-y*\left( \frac{\left| \sum v\left( ATP consumer reactions \right) \right|-minATP}{maxATP-minATP} \right)$$

$$-z*\left( \frac{\left| \sum_{i=1}^{number of reactions} v\left( i \right) \right|-minTotal}{maxTotal-minTotal} \right) \left( 6 \right)$$

The values of minATP, maxATP, minTotal and maxTotal estimated using trial and error and are equal to 0, 10000, 100000, 160000, respectively. We considered y and z in equation 6 equal to one.

There are many different selection methods for the GA [23]. We utilized the stochastic universal sampling method, which is developed based on the fitness proportionate selection method but does not exhibit bias in selection [23].

Also, there are various methods for implementing crossover for GAs, and we applied the single-point crossover method in our implementation. In this method, a random number is generated between one and the total number of traits minus one and call it the crossover location. One child receives traits located before the crossover location from the first parent and traits after the crossover location from the second parent. The second child inherits traits located before the crossover location from second parent and traits after the crossover location from the first parent.

According to the mutation rate in the GA, a predefined number of chromosomes is selected from the population randomly. Then, we picked up a trait randomly and changed its value to another level.

It is suggested that for better performance of the GA, some of the best chromosomes in each generation go to the next generation without any changes in their traits’ levels, which called elitism. We defined the elitism rate using equation 7.

$$Elitism rate=1-crossover rate (7)$$

We designed our GA to stop after a constant number of generations and estimate this constant using trial and error. We set our GA to end iteration at the 100th generation.

To estimate the best values for GA parameters, including population size, crossover rate, and mutation rate, we used the Taguchi method of design of experiments. We define three levels for each parameter and use the L9 table to carry out simulations. The average fitness of individuals at the last generation is considered as response value. As we wanted to maximize fitness values, to calculate the signal to noise ratio we used equation 8.

$$\frac{S}{N}= -10 *log\left( \frac{\Sigma\left( \frac{1}{Y^{2}} \right)}{n} \right) \left( 8 \right)$$

In this formula, S, N, Y, and n are signal, noise, response and number of responses, respectively.

#### Estimation of best population size, crossover rate and mutation rate using the Taguchi method

The initial value for variables such as population size, crossover rate, and mutation rate, have an impact on the GA performance. The L9 Taguchi table, which is used to estimate the suitable GA parameters, and calculated response for each experiment are presented in Table 8.

**Table 8. Design of experiment using Taguchi method and the outcome of each experiment**

| **Experiment number** | **Population size** | **Crossover rate** | **Mutation rate** | **Response** |
| --- | --- | --- | --- | --- |
| 1 | 30 | 0.7 | 0.1 | 5.58 |
| 2 | 30 | 0.8 | 0.2 | 5.19 |
| 3 | 30 | 0.9 | 0.3 | 5.18 |
| 4 | 50 | 0.7 | 0.2 | 5.49 |
| 5 | 50 | 0.8 | 0.3 | 5.16 |
| 6 | 50 | 0.9 | 0.1 | 5.44 |
| 7 | 70 | 0.7 | 0.3 | 5.17 |
| 8 | 70 | 0.8 | 0.1 | 5.59 |
| 9 | 70 | 0.9 | 0.2 | 5.42 |

Using Table 8, we plot our parameter values versus mean of response and signal-to-noise ratio and the result is shown in Fig1.

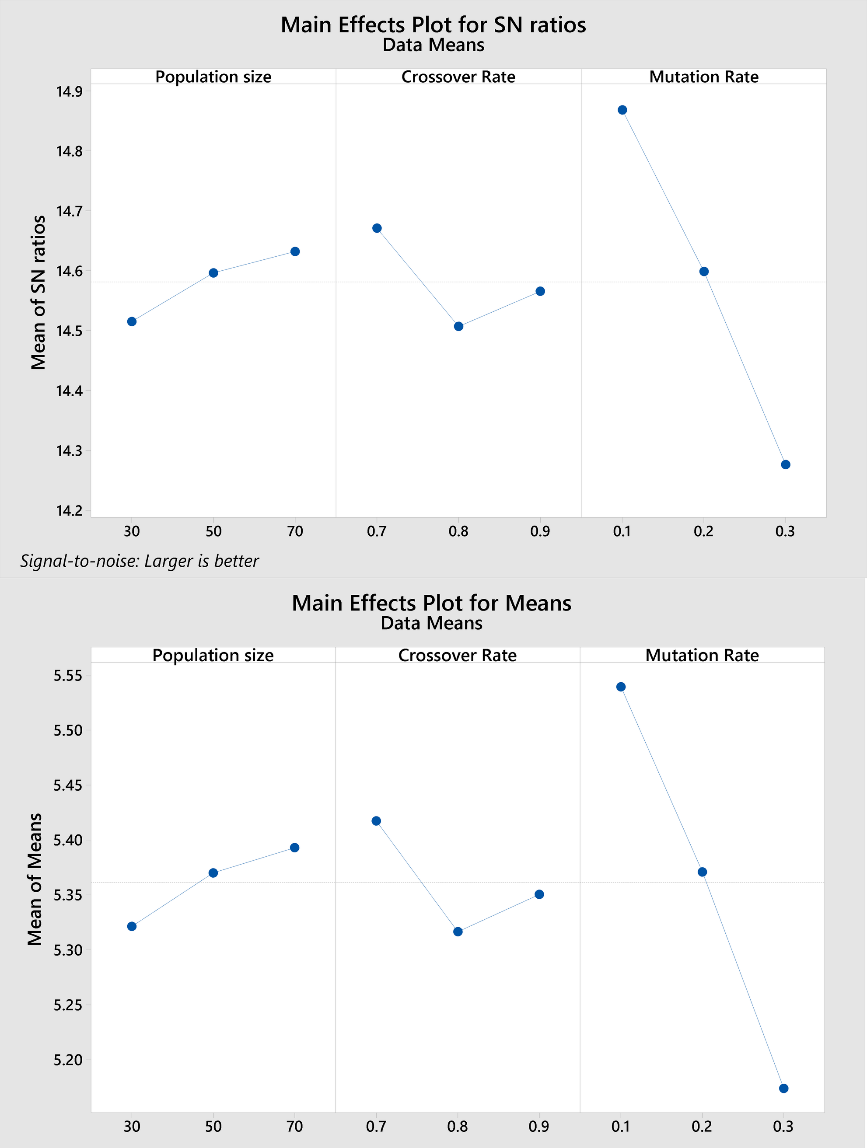

**Fig 1. Visualization of parameters’ effects on the Taguchi design experiments**

According to Fig 1, since the higher the signal-to-noise ratio the better the parameter, we set population size, crossover rate, and mutation rate to 70, 0.7 and 0.1, respectively.

#### Set the limits of scenarios

In the first scenario, all resources were available to the models. In the second scenario, we set the carbon input reaction upper bound to half of its optimal value when the model had the C4 state traits’ level. In the third and fourth scenarios, we use the same method and set the upper bound for nitrogen and water input reaction to simulate nitrogen and water scarcity, respectively. The upper bound for second, third and fourth scenarios are 30, 10 and 60 for carbon, nitrogen, and water input reactions, respectively.

### Visualization of evolutionary events

To visualize the distribution of our individuals on the fitness landscape, we calculated the fitness of all possible individuals in the four scenarios, see the methods section, and then plot the histogram for fitness values in each scenario. The distributions of fitness for all possible individuals in these scenarios are shown in Fig 2. It is important to note that in addition to the distribution of fitnesses, the maximum fitness in various scenarios is different too. These histograms showed us that the fitness landscape varies as we would expect in different scenarios. For example, when there is no limitation on resource availability, as shown in Fig 2 D, the models have higher fitness compared to other scenarios.

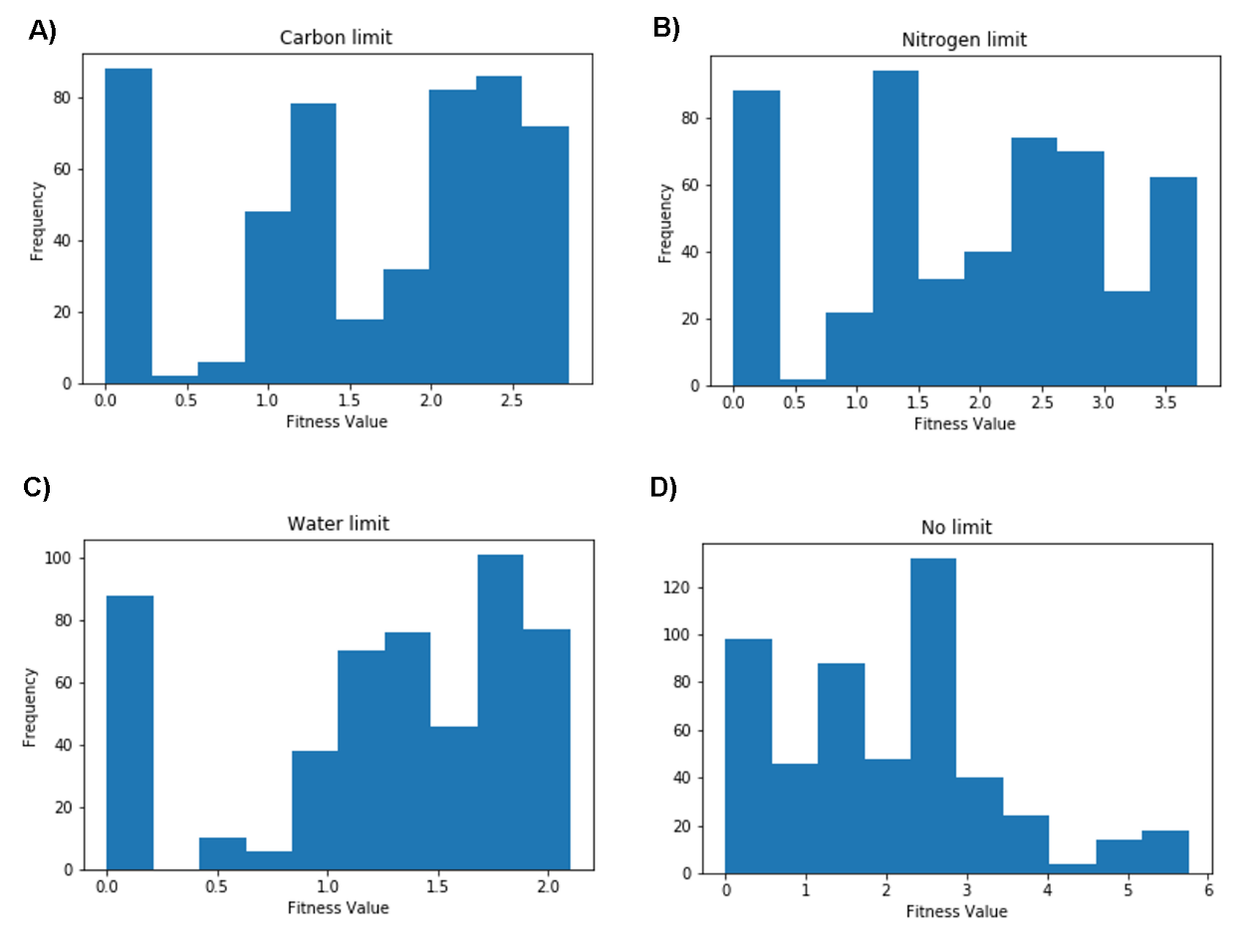

**Fig 2. The fitness value distribution for all possible 512 chromosomes** A) carbon limited scenario B) nitrogen-limited scenario C) water-limited scenario D) No resource limitation scenario

### References

1.	Schlüter U, Weber AP. The road to C4 photosynthesis: evolution of a complex trait via intermediary states. Plant and Cell Physiology. 2016;57(5):881-9.

2.	Mallmann J, Heckmann D, Bräutigam A, Lercher MJ, Weber AP, Westhoff P, et al. The role of photorespiration during the evolution of C4 photosynthesis in the genus Flaveria. Elife. 2014;3:e02478.

3.	Schlueter U, Denton AK, Bräutigam A. Understanding metabolite transport and metabolism in C4 plants through RNA-seq. Current opinion in plant biology. 2016;31:83-90.

4.	Weber AP, Bräutigam A. The role of membrane transport in metabolic engineering of plant primary metabolism. Current opinion in biotechnology. 2013;24(2):256-62.

5.	Scheibe R. Malate valves to balance cellular energy supply. Physiologia plantarum. 2004;120(1):21-6.

6.	Liepman AH, Olsen LJ. Alanine aminotransferase homologs catalyze the glutamate: glyoxylate aminotransferase reaction in peroxisomes of Arabidopsis. Plant Physiology. 2003;131(1):215-27.

7.	Funakoshi M, Sekine M, Katane M, Furuchi T, Yohda M, Yoshikawa T, et al. Cloning and functional characterization of Arabidopsis thaliana d‐amino acid aminotransferase–d‐aspartate behavior during germination. The FEBS journal. 2008;275(6):1188-200.

8.	Linka N, Weber AP. Intracellular metabolite transporters in plants. Molecular plant. 2010;3(1):21-53.

9.	Tronconi MA, Fahnenstich H, Weehler MCG, Andreo CS, Flügge U-I, Drincovich MF, et al. Arabidopsis NAD-malic enzyme functions as a homodimer and heterodimer and has a major impact on nocturnal metabolism. Plant physiology. 2008;146(4):1540-52.

10.	Arnold A, Nikoloski Z. Bottom-up metabolic reconstruction of Arabidopsis thaliana and its application to determining the metabolic costs of enzyme production. Plant physiology. 2014;165(3):1380-91.

11.	Robaina-Estévez S, Daloso DM, Zhang Y, Fernie AR, Nikoloski Z. Resolving the central metabolism of Arabidopsis guard cells. Scientific Reports. 2017;7(1):8307.

12.	Liu L, Shen F, Xin C, Wang Z. Multi‐scale modeling of Arabidopsis thaliana response to different CO2 conditions: From gene expression to metabolic flux. Journal of integrative plant biology. 2016;58(1):2-11.

13.	Sage RF, Khoshravesh R, Sage TL. From proto-Kranz to C4 Kranz: building the bridge to C4 photosynthesis. Journal of experimental botany. 2014;65(13):3341-56.

14.	Gowik U, Bräutigam A, Weber KL, Weber AP, Westhoff P. Evolution of C4 photosynthesis in the genus Flaveria: how many and which genes does it take to make C4? The Plant Cell. 2011;23(6):2087-105.

15.	Edwards EJ. Evolutionary trajectories, accessibility and other metaphors: the case of C4 and CAM photosynthesis. New Phytologist. 2019;223(4):1742-55.

16.	Bräutigam A, Gowik U. Photorespiration connects C3 and C4 photosynthesis. Journal of Experimental Botany. 2016;67(10):2953-62.

17.	Sage RF, Monson RK, Ehleringer JR, Adachi S, Pearcy RW. Some like it hot: The physiological ecology of C 4 plant evolution. Oecologia. 2018;187(4):941-66.

18.	Leegood RC. Roles of the bundle sheath cells in leaves of C3 plants. Journal of Experimental Botany. 2008;59(7):1663-73.

19.	Bräutigam A, Kajala K, Wullenweber J, Sommer M, Gagneul D, Weber KL, et al. An mRNA blueprint for C4 photosynthesis derived from comparative transcriptomics of closely related C3 and C4 species. Plant physiology. 2011;155(1):142-56.

20.	Schlüter U, Bräutigam A, Gowik U, Melzer M, Christin P-A, Kurz S, et al. Photosynthesis in C3–C4 intermediate Moricandia species. Journal of Experimental Botany. 2016;68(2):191-206.

21.	Aubry S, Brown NJ, Hibberd JM. The role of proteins in C3 plants prior to their recruitment into the C4 pathway. Journal of experimental botany. 2011;62(9):3049-59.

22.	Reeves G, Grangé-Guermente MJ, Hibberd JM. Regulatory gateways for cell-specific gene expression in C4 leaves with Kranz anatomy. Journal of experimental botany. 2017;68(2):107-16.

23.	Baker JE, editor Reducing bias and inefficiency in the selection algorithm. Proceedings of the second international conference on genetic algorithms; 1987.
